# A coupled mechano-biochemical framework for root meristem morphogenesis

**DOI:** 10.1101/2021.01.27.428294

**Authors:** Marco Marconi, Marçal Gallemi, Eva Benková, Krzysztof Wabnik

**Author notes:** **Author Contributions**: KW and MM designed the computer model and analyzed data. M.M implement model and performed computer model simulations. MM and KW wrote the manuscript with the help from co-authors. MG and EB design and performed time-lapse confocal imaging experiments. **Competing Interest Statement**: the authors declare no competing interest.

## Abstract

Plants grow roots to adjust their bodies to dynamic changes in the surrounding environment. How do plant roots emerge from the stem cell reservoir during embryogenesis is however poorly understood. Here, we present a bottom-up strategy to address this challenge by combining empirical observations with advanced computer modeling techniques. We demonstrate that the anisotropy of root growth results from differential growth rates of adjacent tissues, whereas the root meristem development incorporates a multi-level feedback loop between complex transport network of phytohormone auxin, auxin-dependent cell growth and cytoskeleton rearrangements. *In silico* model predictions are in close agreement with *in vivo* patterns of anisotropic growth, auxin distribution, and cell polarity, as well as several root phenotypes caused by chemical, mechanical, or genetic perturbations. Our findings reveal a minimal set of design principles connecting tissue mechanics, cell anisotropy, and directional transport that are sufficient for self-organization of the root meristem shape. A mobile auxin signal transported through immobile cells orchestrates polarity and growth mechanics to instruct the morphogenesis of an independent organ.

## Introduction

Self-organization is a fundamental attribute of life that is present at multiple scales, from complex ecosystems to individual atoms in proteins. In that sense, plants are remarkable self-organizers as they successively produce organs *de novo* from stem cell reservoirs to adapt to the dynamic changes in the environment. Despite the lack of a central nervous system, organs such as roots can coordinate their growth through local interactions between adjacent non-mobile cells to sustain the optimal exploration of soil-derived resources(O’Brien et al., 2016), and provide water and mineral absorption as well as a stable anchor on the ground(Chapman et al., 2012). The root of model plant *Arabidopsis thaliana* shows clear signs of anisotropic growth during late stages of embryogenesis(Daher et al., 2018). Root meristem maturation is then followed by a sequence of asymmetric cell divisions and cell expansion that involve the phytohormone auxin(Adamowski and Friml, 2015) and associated changes of cytoskeleton components such as cortical microtubules (CMTs)(Adamowski et al., 2019; Siegrist and Doe, 2007). Whereas CMTs may respond to mechanical stresses(Hamant et al., 2019; Hamant and Haswell, 2017) by constraining the directional trafficking of membrane proteins, small signaling molecules and cell wall elements(Adamowski et al., 2019; Siegrist and Doe, 2007), auxin indirectly controls the weakening of the cell walls, promoting cell elongation(Barbez et al., 2017; Majda and Robert, 2018).

Unlike any other phytohormone, auxin is transported in a directional (polar) manner(Wisniewska et al., 2006). The direction of auxin transport is associated with the polar subcellular localization of the PIN-FORMED (PIN) auxin efflux carriers(Adamowski and Friml, 2015), which determines auxin allocation in the root to instruct the cell growth(Benková et al., 2003). This apparent complexity of root growth and patterning mechanisms has long attracted theoreticians on the quest to identify the underlying principles. In the last decade, several computational models of root development yielded important insights about auxin-dependent growth and zonation of mature roots(Morales-Tapia and Ruz-Ramírez, 2016; Rutten and ten Tusscher, 2019; Wabnik et al., 2011). However, these root models typically incorporate pre-defined patterns of polar auxin flow on idealized geometries (e.g. cell grids)(Band et al., 2014; Grieneisen et al., 2007; Mähönen et al., 2014; Mironova et al., 2010; Wabnik et al., 2013) and rarely integrate growth mechanics(De Vos et al., 2014; Fozard et al., 2013; Jensen and Fozard, 2015). The major challenge is, however, to identify the so far elusive mechanisms that generate initial symmetry breaking leading to anisotropic root growth and establishment of a sophisticated polar auxin transport network. A robust approach to this challenge must accommodate both biochemical and biomechanical aspects of early organ growth patterns which are equally challenging to develop(Delile et al., 2017; Fletcher et al., 2014; Heisler et al., 2010).

Here, we address mechanisms of symmetry breaking in the root meristem morphogenesis using a bottom-up computer modelling strategy. Our approach organizes empirical data into plausible mechanisms for the establishment of auxin-dependent root growth, tissue biomechanics, and polar auxin transport network. Furthermore, our model of the root meristem operates at single cell resolution, and does not require polarity pre-patterning to predict growth anisotropy and auxin distribution observed *in vivo*. Through model simulations we gather mechanistic insights in the *de-novo* establishment of cell polarities and root growth anisotropy from an initially non-polar scenario. Our study suggests that a root meristem organization can be predicted from local interactions between directional transport of auxin, auxin-dependent cell elongation, cell polarization, and biomechanical stimuli.

## Results

### Anisotropic root growth results from differential expansion at the root-shoot interface

Plant embryogenesis follows the fertilization of the egg, and successive formation of the zygote(Park and Harada, 2008). The zygote quickly elongates and sets the embryonic axis within a few hours, and after several cell divisions the aerial and root parts are already clearly distinguishable(Kimata et al., 2016). The stem cell niche is initiated during the heart developmental stage(Ten Hove et al., 2015). Therefore, we chose this stage as the starting point for the construction of the computer model of the emerging root meristem. By digitizing the confocal microscopy images of the heart-stage *A. thaliana* embryo we created 2D meshes with MorphoGraphX(de Reuille et al., 2015) (Supplementary Fig. S1). Next, we developed a new biomechanical growth framework using the Position-based Dynamics (PBD) techniques(Müller et al., 2007) (Supplementary Fig. S2A). PBD is a fast, stable and controllable framework(Müller et al., 2007) ideally suited for subcellular resolution models and thus presents an efficient alternative to computationally demanding continuum mechanics approaches such as Finite Element Methods(Bidhendi and Geitmann, 2018; Fayant et al., 2010). In our framework the growth mechanics are defined by a set of constraints that include internal turgor pressure, viscoelastic behavior of plant cell walls and mechanical deformations (strain) (Supplementary Fig. S2B and Supplementary Information for model details).

Next, we explored through our model the potential origin of anisotropic root growth. Recent studies suggest that differential cell growth produced by mechanical interactions may regulate organogenesis independently from genetic control, and potentially feedback on it(Hervieux et al., 2017; Kierzkowski et al., 2012). We hypothesized that differential growth rates at the root-shoot interface (RSI) during embryogenesis could potentially trigger this initial symmetry breaking (Fig. 1A-B). We performed model simulations by assuming uniform growth of root and RSI (Fig. 1A, Supplementary Video S1). In this scenario, we could only observe the strong isotropic growth of the root radicle without a specific orientation of CMTs (Fig. 1A). Typically CMTs are orthogonal to the principal strain direction(Hamant et al., 2019), however, the lack of any mechanical growth restriction eventually led to a purely isotropic deformation. In contrast, by assuming a faster growth of the radicle compared to the adjacent embryonic tissues results in a strong anisotropy expansion with clearly visible root polarity axis (Fig. 1B and Supplementary Video S1). The likely reason for this observation is that a slowly growing RSI prevents the expansion of the faster growing radicle in the radial direction; the radicle is therefore forced to enlarge longitudinally and the resulting strain allows the CMTs to reorient themselves and feedback on the radicle growth creating the desired anisotropy. We tested these model predictions by quantifying the growth increase over a 6 hour period in radicle and hypocotyl of young seedling using time-lapse confocal microscopy imaging (Fig. 1C)(Zhu et al., 2019). Indeed, the emerging root radicle grew ∼4x faster than the adjacent hypocotyl (Fig. 1D) which supports our hypothesis.

**Fig. 1.**
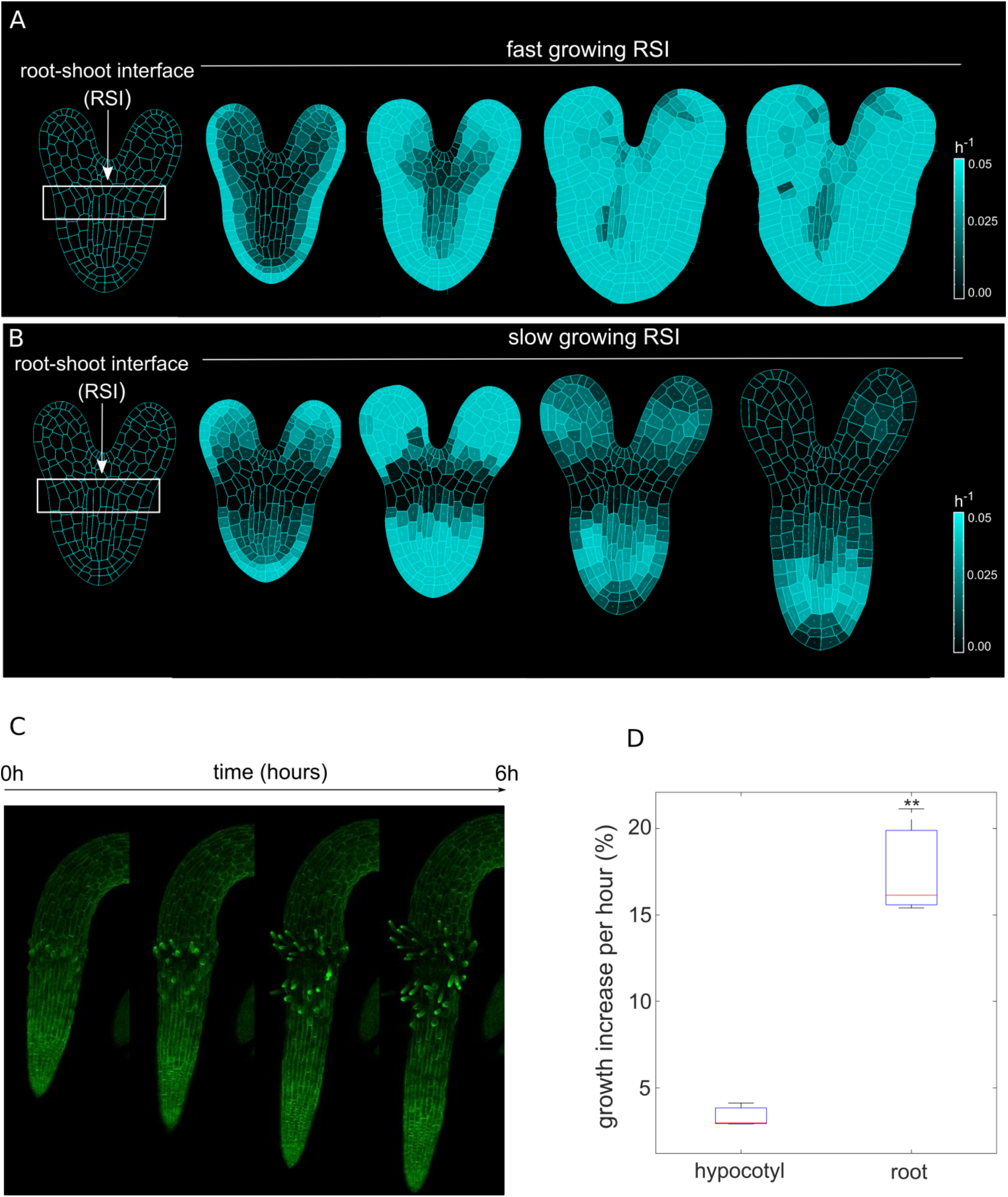
Differential cell growth at the embryo RSI produces anisotropic expansion of the root. (**A**) Simulated root embryo with uniform growth rate between the root and the RSI. If the RSI and the root are allowed to growth at the same rate, the root is unable to achieve independent anisotropic growth since there is no clear strain direction acting on the root cells which could allow CMTs re-orientation. As a consequence, the whole embryo expands isotropically. (**B)** Simulated root outgrowth by assuming differential growth rate between the RSI and the root. Under these condition, the root tissues are unable to expand radially because of the slower growing RSI; the root is therefore forced to expand longitudinally and the resulting strain allow CMTs re-orientation orthogonal to the strain direction, making anisotropic growth possible (**C**) Screenshots from time-lapse imaging of growing radicle with PIP-GFP plasma membrane marker(Zhu et al., 2019) for a total time of 6 hours. (**D**) Percentage of growth increase per hour for adjacent organs radicle (root) and hypocotyl quantified from the time-lapse confocal imaging of three independent plants (n=3). **p-value < 0.015 a one-way ANOVA.

In summary, model simulations and experiments jointly suggest that the anisotropic growth of the root could emerge from initially uniform isotropic cell expansion guided by differential growth rates at the root-shoot interface. This oriented growth further restricts root elongation, primary along the longitudinal axis.

### Feedback between anisotropic growth and auxin flow shapes *Arabidopsis* root pattern

Our model suggests that differential growth at the RSI could trigger non-uniform CTMs orientation and determine the root growth axis. In *A. thaliana* root, cell growth rates are precisely regulated by cellular auxin levels that can be tuned through intercellular transport, involving auxin influx and efflux carriers(Adamowski and Friml, 2015). While auxin influx carriers of the AUX1/LAX family are typically uniformly distributed around the cell membranes(Kleine-Vehn and Friml, 2008), PIN auxin efflux carriers show polar subcellular localization in the root that direct the auxin flow in (rootward) or out (shootward)(Wisniewska et al., 2006). However, mechanisms underlying PIN polarization establishment in the root are unclear(Feraru et al., 2011). CTMs restrict the deposition of various protein cargoes on the plasma membrane, typically along the maximal growth direction (maximal strain) (Adamowski et al., 2019; Nieuwland et al., 2016; Siegrist and Doe, 2007; Yang, 2008). It is plausible that PIN protein allocation (and/or other cargoes) at the plasma membrane might be restricted by CTMs. However, CMTs would only restrict the axis along which PIN trafficking will occur, and thus CMTs alone cannot determine the exact direction (rootward or shootward) in which the auxin would flow.

Auxin affects trafficking of PIN proteins by an unknown mechanism. Several theories of how PIN polarities are established have been proposed, i.e. through sensing the net auxin flux through the cell(Feugier and Iwasa, 2006; Mitchison, 1980; Rolland-Lagan and Prusinkiewicz, 2005; Stoma et al., 2008), auxin concentrations(Jönsson et al., 2006; Merks et al., 2007; Smith et al., 2006; Wabnik et al., 2010), the auxin gradient inside the cell(Kramer, 2009) or a combination of those(Cieslak et al., 2015). It is also known that auxin affects endocytosis of PINs by modulating their transport(Narasimhan et al., 2021). We propose two scenarios that are compatible with this experimental observation. In theory cells could sense auxin flux through the membrane (also called “with-the-flux model”)(Feugier and Iwasa, 2006; Mitchison, 1980; Rolland-Lagan and Prusinkiewicz, 2005; Stoma et al., 2008) and adjust PIN allocation to the plasma membarne in a feedback dependent manner (Supplementary Fig. S3A-B). The molecular mechanism behind auxin flux sensing is unclear but could involve membrane-associated protein kinases(Hajný et al., 2020; Marhava et al., 2018; Michniewicz et al., 2007). Alternatively, we propose a second molecular mechanism for PIN polarization compatible with experimental observations(Narasimhan et al., 2021) which we refer to as ‘regulator-polarizer’ (Supplementary Fig. S3C-D). In this model, auxin passes through a plasma membrane and activates a putative regulator (i.e. general phosphatase(Michniewicz et al., 2007)) that is allowed to freely diffuse in the plasma membrane. The diffusing regulator inhibits the local polarizer that activates PINs (e.g. dedicated protein kinase that phosphorylates PIN(Hajný et al., 2020; Michniewicz et al., 2007)) and controls PINs deposition on the membrane. This concept is similar to a classical idea of reaction-diffusion systems(Green and Sharpe, 2015). The current model does not distinguish between different PIN families(Sauer and Kleine-Vehn, 2019), instead ‘general’ PINs are distributed according to this unified mechanisms of polarization (see SI for more details). Regarding auxin importers, the model only implements AUX1/LAX carriers(Swarup et al., 2001) by assuming uniform distribution over the cell membrane, while other known importers like ABCB transporters(Cho and Cho, 2013) are not currently considered.

Previous modeling attempts combined auxin transport with tissue mechanics to explain a unidirectional PIN polarity pattern associated with the shoot apical meristem but operated on static non-growing templates(Heisler et al., 2010). We therefore put together growth biomechanics (Fig. 1) and polar auxin transport models into a coherent mechanistic framework (Supplementary Fig. S4, Supplementary Fig. S5), and tested whether this framework is capable of generating complex PIN polarity network. This network would guide the auxin-dependent root growth mechanics to control the root meristem shape. Computer model simulations follow the growth of an embryo radicle (immature root) that is connected to the aerial segment of the plant embryo (Fig. 2A). Auxin is introduced into the vascular tissues and exit through the epidermis (Fig. 2A, Supplementary Fig. S4B), allowing the auxin recycling between the emerging root and the rest of the embryo(Möller and Weijers, 2009). In the simulations, internal turgor pressure balances the cell walls stiffness, and auxin can disrupt this balance by reducing the stiffness of the cell walls, thereby promoting cell wall elongation(Majda and Robert, 2018)(Fig. 2B, Supplementary Fig. S4C). To incorporate quantitative data in the model, parameters were fit to the experimental kinetics of PIN trafficking after cell division (Fig. 2K)(Glanc et al., 2018). In our model cell division follows a simple but effective rule: each cell possesses a maximum area attribute, so that when a cell reaches a certain threshold it divides in two daughter cells (Supplementary Fig. S4D). The maximum area is specific for each cell type, so that cell size is maintained consistent for each cell lineage. The orientation of cell division is by default anticlinal and occurs along a division vector passing through the cell centroid and perpendicular with the growth direction (Supplementary Fig. S4D).

**Fig. 2.**
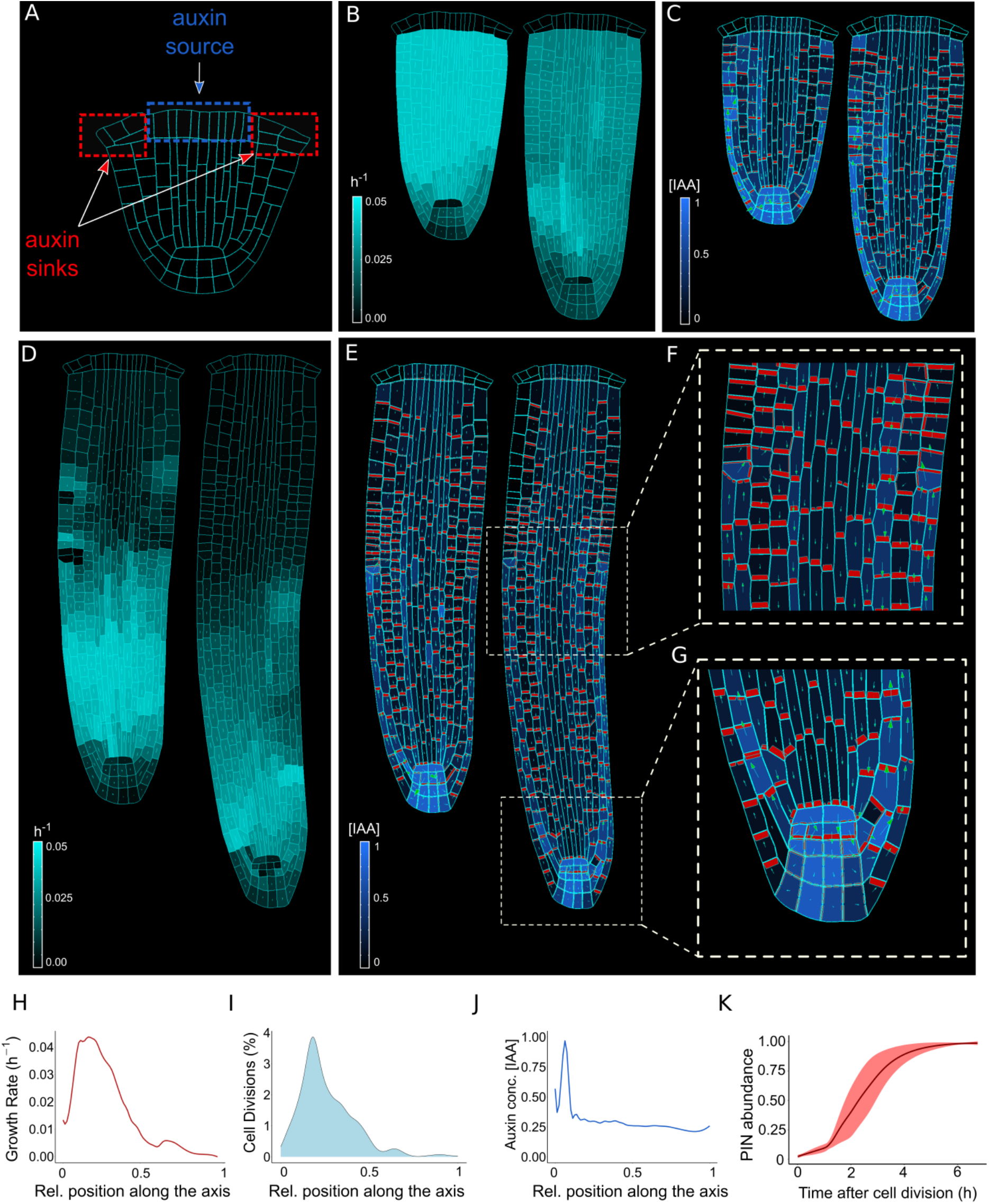
The model reproduces realistic root meristem geometry, auxin distribution and PINs polar localizations. (**A**) Initial embryonic set point. Locations of auxin influx (auxin source, blue) and evacuation (auxin sinks, red) from the embryo are shown. (**B-E**) Model simulations predict a time evolution of cell growth rates (B and D, bright cyan color) and principal growth directions (white lines). C, E dynamics of auxin distribution (blue color), auxin flow direction (arrows) and PIN localizations (red). Ongoing cell division events are marked by black regions. (**F, G**) Zoom on basal meristem (F) and root apical meristem (G). The model correctly reproduces very detailed PINs localizations including bipolar PIN2 localization in the cortex (F). (**H**) Growth rate profile along the root axis. The fastest growing region is located in the apical meristem as observed experimentally(Bassel et al., 2014). (**I**) Cell division frequencies along the root axis. The majority of cell divisions occur in the apical meristem. (**J**) Auxin gradient along the root axis. Most of auxin is concentrated in the root tip as observed in experiments(Overvoorde et al., 2010). (**K**) Time lapse profile of PINs re-localization on the membranes after cell division event. PINs re-localization is completed in approximately 5-6 hours after cell division(Glanc et al., 2018).

Time-lapse model simulations predicted the anisotropic auxin-dependent root growth (Fig. 2B and 2D) with a peak located in the apical root meristem (Fig. 2H and 2I). This prediction is in a close agreement with experimental observations(see Fig. 2 in (Bassel et al., 2014)). Furthermore, growth anisotropy (Fig. 2D and Supplementary Fig. S6) and associated cell division patterns (Fig. 2I) follow the predicted auxin distributions (Fig. 2C, 2E, and 2J), eventually reproducing the complex shape of the root (Fig. 2G, Supplementary Fig. S7, Supplementary Videos S2 and S3), regardless of choice of the polar auxin transport scenario(Supplementary Fig. S4E). The predicted auxin maximum formed close to the Quiescent Center (QC) (Fig. 2G) and represents the equilibrium between auxin reaching the root tip from the vascular tissues and auxin leaving the root tip to the outermost tissues. Previous evidence showed that this auxin maximum is necessary for the correct organization of the meristem(Petersson et al., 2009). However, to obtain a realistic shape of root tip additional assumptions was necessary considering the cells belonging to the stem cell niche, which are known to follow alternative division rules(Fisher and Sozzani, 2016); cortex/endodermis initial daughters (CEID) and the epidermis/lateral root cap initials divide periclinal and alternatively anticlinal/periclinal, respectively, as previously described(Choi and Lim, 2016). The removal of this assumption from the model produced an incorrect pattern of cell divisions in ground tissues and slightly altered root morphology (Supplementary Fig. S8).

Lastly, our biochemical-mechanical root model was able to predict a complex PIN polarization network from an initially non-polar scenario (Fig. 2E-G, Supplementary Fig. S7E-G and Supplementary Video S2 and S3). This dynamic network includes rootward PIN localization in vascular tissues and shootward localization in outermost epidermis that closely follow experimentally observed patterns (see Fig. 1 in (Billou et al., 2005) and Fig. 2 in (Tanaka et al., 2006)). We also tested the model robustness to cellular geometry and key model parameters that control PIN polarity and auxin effect on cell growth. Choosing alternative wild-type embryo geometry(Nieuwland et al., 2016; Scheres et al., 1994) has no visible impact on root anisotropy, auxin distribution and PIN patterns in the simulations (Supplementary Fig. S9A-B). Similarly, model predictions were robust in a plausible range of parameter values (Supplementary Fig. S10A-B).

An intriguing prediction of the model was the bidirectional (shootward and rootward) localization of PIN proteins in the cortex tissues (Fig. 2F and Supplementary Fig. S7F) in the transition region that corresponds to the end of the lateral root cap (LRC). This exact ‘bipolar’ PIN localization in the cortex has been previously observed in experiments close to the transition zone(Ötvös et al., 2021; Sauer et al., 2006), but the function of this phenomenon remains unknown. We suggest that bipolar PIN localization in the cortex is the likely the result of the bidirectional auxin flux from LRC and epidermis into cortex that conflicts the rootward auxin flow in vascular tissues and shootward auxin flow in epidermis (Fig. 2F and Supplementary Fig. S7F). In the next section we will discuss predicted function of bipolar PIN localization for root patterning.

Taken together, computer simulations indicate that our model framework is sufficient for the transformation of the embryonic radicle into a mature root meristem.

### Shoot-independent root growth requires auxin reflux, local auxin production, and balance in auxin levels

Our model simulations predict the self-organization of PIN auxin transport network (Fig. 2F-G, Supplementary Fig. S7F-G), suggesting the presence of lateral auxin transport from the external tissues (epidermis and LRC) into cortex and the stele (Fig. 2E, Supplementary Fig. S7E). Predicted ‘bipolar’ PIN localization (Fig. 2F, Supplementary Fig. S7F) could drive polar auxin redistribution towards inner tissues, which fuels a phenomenon called reflux loop(Benková et al., 2003; Grieneisen et al., 2007; Paponov et al., 2005). This lateral auxin transport between epidermis and cortex might be supported by plasmodesmata-dependent diffusion(Mellor et al., 2020) but its directionality is critical for the growth coordination of adjacent epidermis and cortex tissues(Ötvös et al., 2021). However, it is unknown how this reflux phenomenon would be retained in realistic tissue geometries under mechanical constrains(Grieneisen et al., 2007).

To further investigate the importance of the dynamic PIN localization network for the sustained growth of the root, we performed model simulations by preventing lateral auxin transport (Fig. 3A, B). We found that a negligible amount of auxin was able to enter the cortex, as no lateral auxin influx from the epidermis was present, and bipolar PIN localizations was lost in ‘no-reflux’ simulations (Fig. 3C and Supplementary Video S4) compared to the reference reflux scenario (Fig. 3D and Supplementary Video S4). This prediction confirms the importance of PIN-mediated lateral transport in cortex for auxin redistribution in the inner root tissues. Despite this, the lack of auxin recycling in the meristem has no evident outcome on root growth rates as long as a reliable auxin supply from the shoot was present (Fig. 3E). Therefore, we also separated the root from the rest of the plant by removing the shoot-derived auxin source (Fig. 3A-B, bottom panel). In the scenario with no-reflux and bipolar PIN localization, root growth could not be sustained over prolonged time (Fig. 3E) and the auxin content inside the root eventually decayed unless an additional auxin source in the root was present (Fig. 3F). In contrary, the reflux scenario allows for the maintenance of auxin levels over prolonged period of time even without shoot derived auxin. Root growth can be further strengthen by incorporating an auxin biosynthesis in the QC cells(Stepanova et al., 2008), which is able to sustain root growth almost indefinitely(Fig. 3E-F). These results together indicate that the presence of a auxin reflux in mediated by bidirectional PIN transport and diffusion in cortex/epidermis can promote the root growth independence for prolonged periods.

**Fig. 3.**
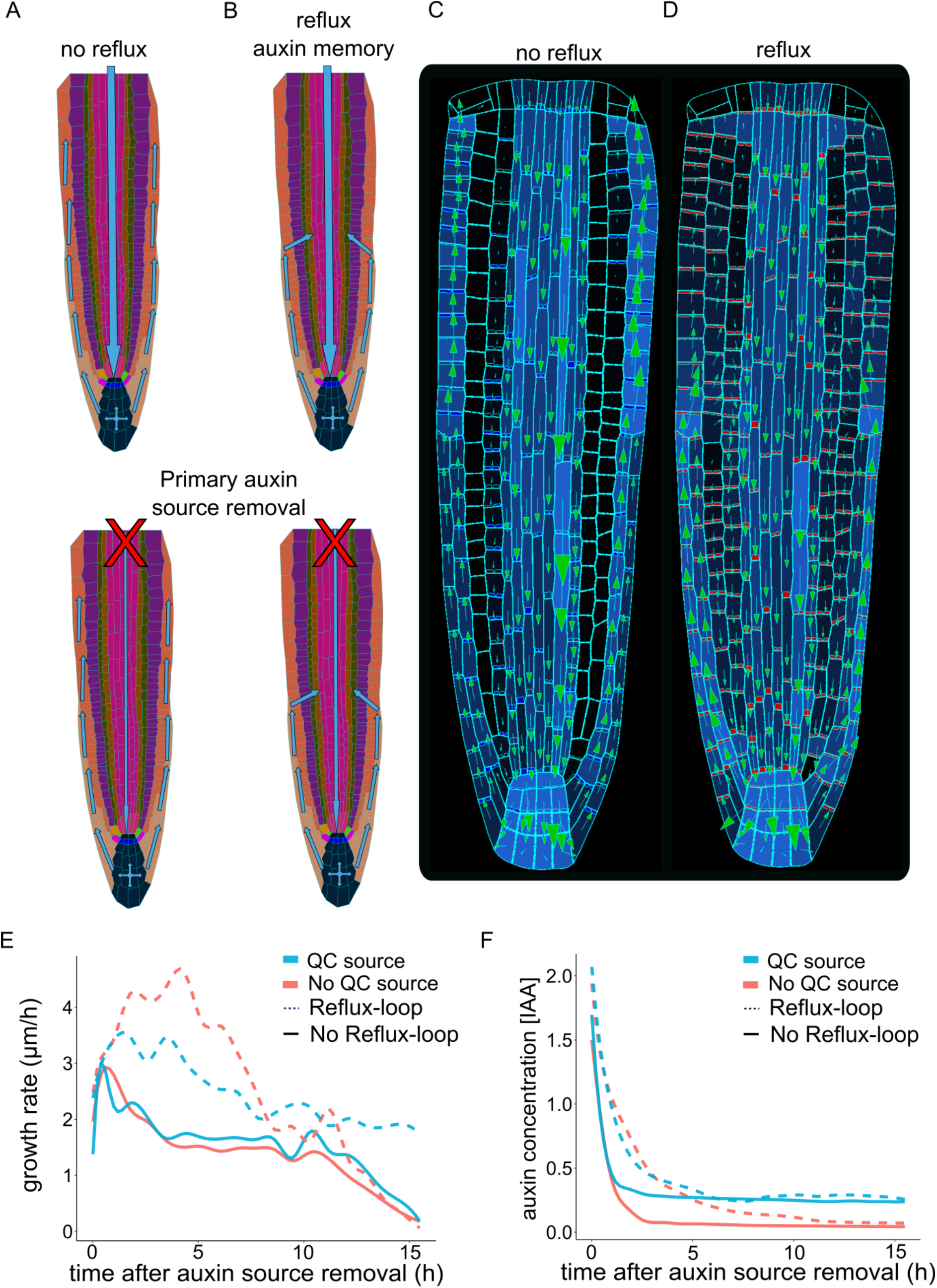
Independent root growth requires auxin reflux and local auxin production. (**A-D**) Schematics (A and B) and model simulations (C and D) with the disabled auxin reflux-loop (A, C) or wild-type-like scenario with self-emerging reflux (B, D). Only in reflux scenario auxin is permitted to lateralize from the epidermis back into the vascular tissues sustaining the long-term root growth. (**E**) Growth rate profiles of model simulations after primary auxin source removal, in four different scenarios. In the absence of an auxin reflux-loop the root is unable to sustain growth for a long period of time (solid lines) even if a secondary auxin source in the root tip was introduced (solid blue line). On the contrary, the presence of an auxin reflux-loop allows the root to sustain growth for prolonged periods of time (dotted lines), even further augmented by the presence of a secondary auxin source in the root tip (dotted blue line). (**F**) Auxin concentration profiles of model simulations after primary auxin source removal, under different scenarios. In the absence of an auxin reflux-loop the average auxin concentration in the root quickly drops to zero (solid red line). Alternatively, the presence of an auxin reflux-loop allows the root to maintain its auxin reserve for prolonged periods of time (dotted blue line). The presence of a secondary auxin source in the root tip allows the root to further preserve its auxin reservoir for extended period (blue lines).

Maintaining the correct balance in auxin levels may also be important for sustaining root growth mechanics in addition to auxin reflux and auxin production. We next tested how alterations in auxin levels alone would impact on the mechanics of root growth dynamics. For that purpose, we successively simulated external auxin applications every 6 hours by increasing overall auxin content of the root (Fig. 4A-B and Supplementary Video S5). Model simulations predict the sequential inhibition and reinstatement of root growth after cyclic auxin removal (Fig. 4C). A similar trend was observed for a shorter period of stimulation (Supplementary Fig. S11). Notably, these model predictions closely agree with the experimentally-observed temporal inhibition of root growth by external auxin applications (see Fig.1f in (Fendrych et al., 2018)).

**Fig. 4.**
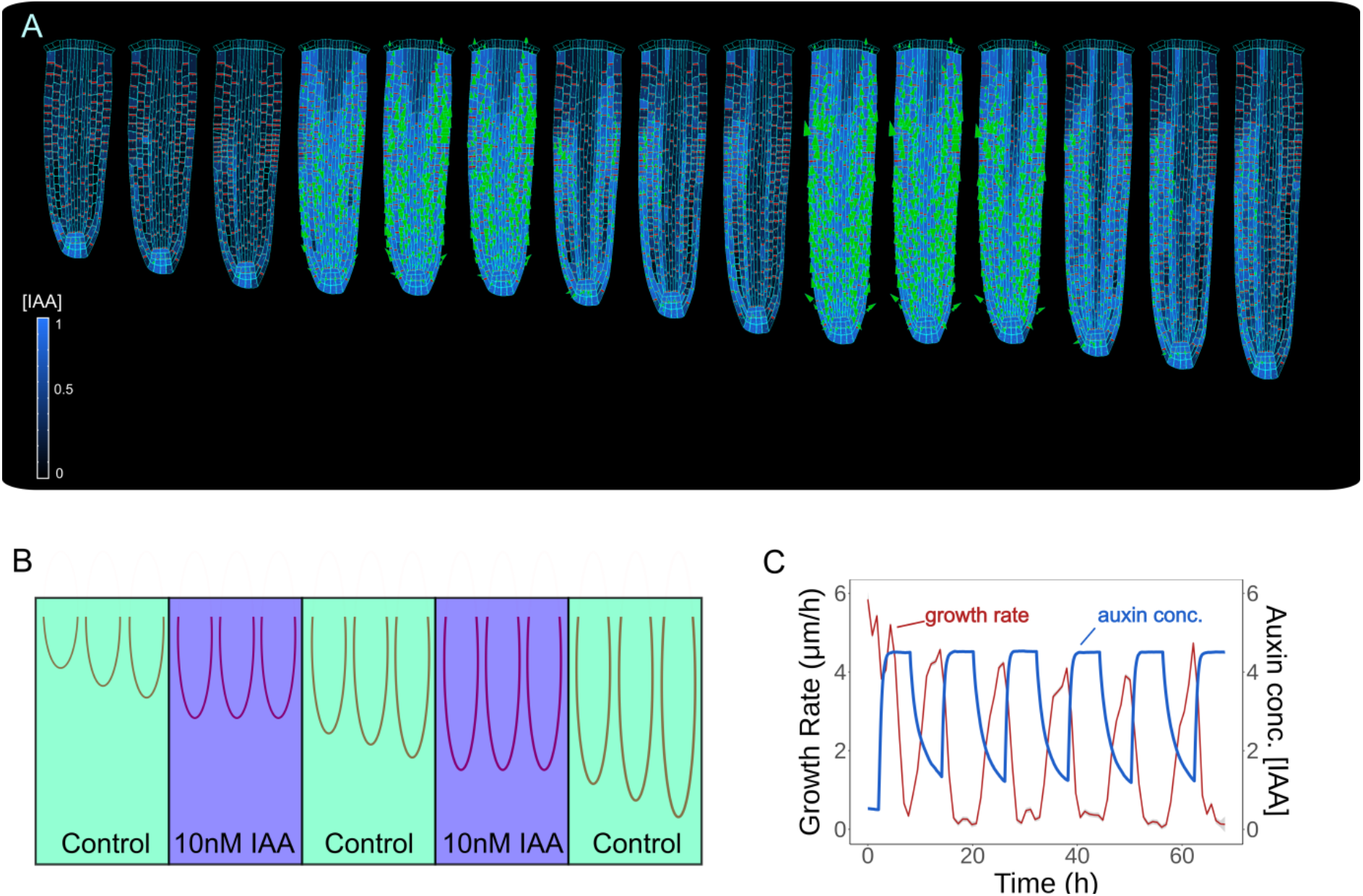
Model predictions successfully reproduce auxin-dependent reversible inhibition of root growth. (**A**) Successive application of external auxin in model simulations according to a predefined cycle. Root growth is inhibited by the introduction of high amounts of auxin, and subsequently restored after the external application is stopped as seen in experiments(Fendrych et al., 2018). (**B**) Schematic of the *in silico* experiment. To simulate auxin treatment as described in(Fendrych et al., 2018), we uniformly introduced external auxin inside the root (by inducing auxin synthesis at cellular level) at predefined time points to inhibit root growth and subsequently removed to allow root growth re-establishment. (**C**) Time lapse profile of root growth rate (red line) and average cell auxin concentration (blue line) monitored during *in silico* experiment. Introduction of external auxin inhibits root growth rate which is subsequently restored after external auxin is removed.

Our analysis indicates that the root model can correctly recapitulate experimentally observed oscillatory root growth response to periodically applied auxin. The model suggests that the self-sustained balance in auxin content of the root mediated by emerging PIN polarity network is critical for the sustained meristem growth mechanics.

### Model simulations predicts experimentally observed phenotypes of auxin-mediated growth and mechanical stress perturbations

Our analysis combined with experimentally validation of model predictions indicates that our mechanic-biochemical model of root meristem morphogenesis could be potentially used to test dynamic perturbations, such as genetic alterations and mechanical manipulations, further focusing design of experiments in the laboratory. To test the predictive power of our model we investigated how alterations of auxin transport parameters could perturb patterning dynamics and whether those predictions would match experimental observations.

PIN2 is an important auxin efflux carrier expressed in the most external root tissues: cortex, epidermis and lateral root cap(Adamowski and Friml, 2015), and steers root gravitropic responses(Rahman et al., 2010). PIN2 loss-of-function results in defective gravitropic response largely because of disrupted auxin transport dynamics(Dhonukshe et al., 2010). To test whether our model could predict the alterations of auxin-distribution phenotype of *pin2* knockdown mutant, we performed computer simulations by reducing PIN expression rate in epidermis, cortex and lateral root cap (Fig. 5A-B and Supplementary Video S6). The reduced levels of PIN in these tissues resulted in auxin accumulation in the lateral root cap on both sides of the root (Fig. 5B), something absent from the wild-type simulations (Fig. 5A). These predictions mimic experimental observations of PIN2 knockdown mutant (see Fig. 2F in (Liu et al., 2018)). Similarly, reduced expression of PIN transport in the vascular tissues resulted in severe alteration of auxin distribution and growth defects(Vieten et al., 2005) (Supplementary Fig. S12A and Supplementary Video S7). Finally, we tested how a general knockdown of auxin cellular influx would impact on root growth. Severely reducing auxin influx by 90% led to lower auxin content, reduced sensitivity to auxin and thereby slow root growth (Supplementary Fig. S12B and Supplementary Video S8) as previously suggested(Inoue et al., 2016; Liu et al., 2018).

**Fig. 5.**
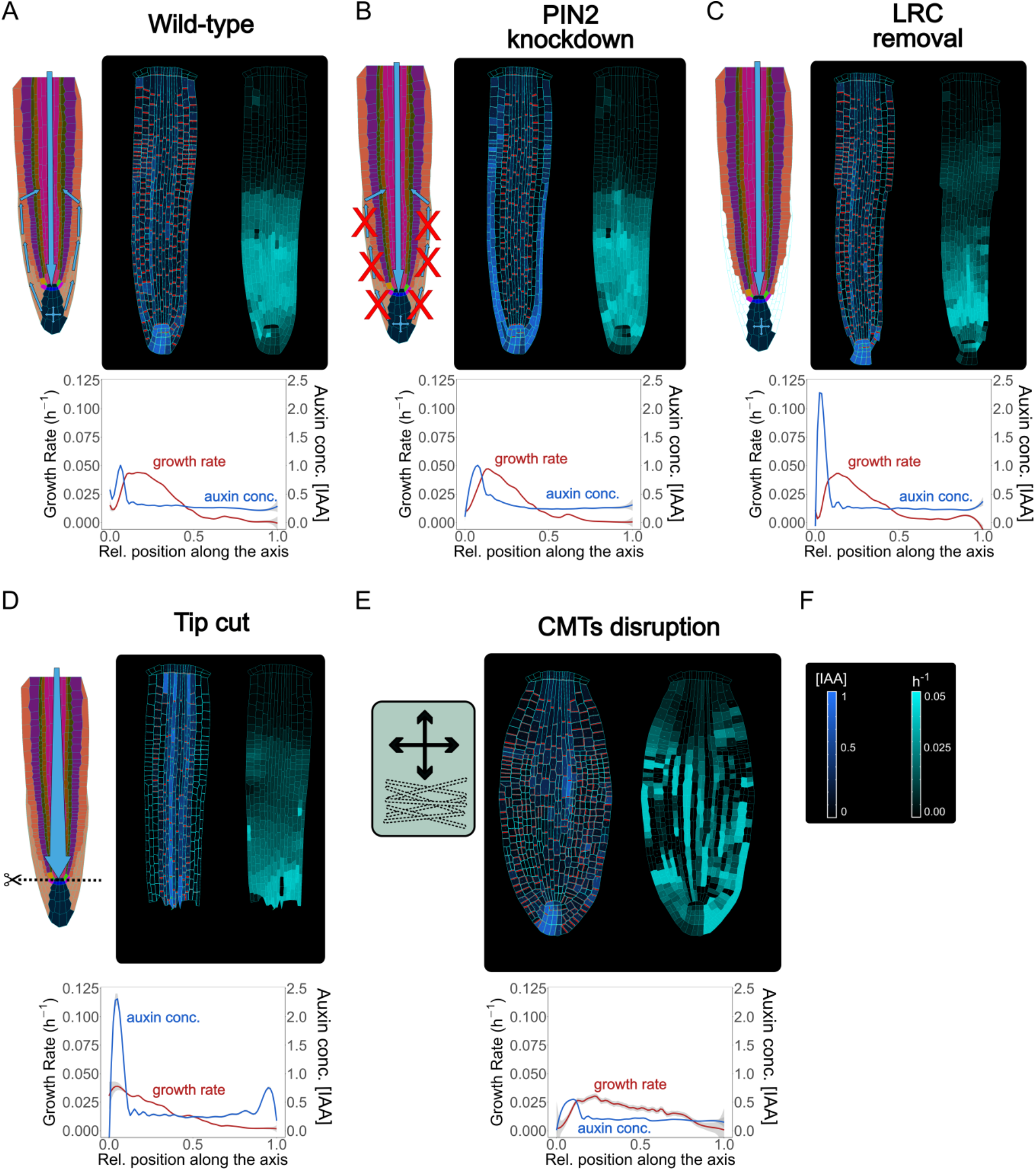
Model simulations recapitulate experimentally observed phenotypes through genetic, pharmacological and mechanical perturbations. (**A**) Reference model simulation of the wild-type scenario. The figure displays a schematic representation of the auxin flow inside the root (left), auxin distribution and cell growth rate from the model simulations (right), and profiles of auxin concentration (blue line) and growth rate (red line) along the root axis (bottom). (**B**) Model simulation of the *pin2* knockdown mutant. *In silico pin2* mutant shows strongly reduced PINs expression in the lateral root cap, epidermis and cortex. Note that acropetal auxin flow is severely affected and auxin tends to accumulate in the lateral tissues as observed in experiments(Dhonukshe et al., 2010). (**C**) Mechanical removal of lateral root cap resulted in the strong accumulation of auxin inside the root tip, largely because auxin cannot flow anymore shootwards through outermost tissues whereas growth rate was not significantly affected. (**D**) Simulation of root tip cutting. Removing the root tip results in a general increase of auxin level in the central vascular tissues, as a result of the disappearance of acropetal auxin flow. Growth rates profiles demonstrate a significant reduction of root growth by nearly 50%. (**E**) Simulated CMTs disruption (i.e. oryzalin treatment or similar) on root growth and polarity. CMTs disruption was simulated by inducing a fast degradation of cortical microtubules. Cells lose polarity and growth anisotropy, causing the root to expand and bulge radially as observed in experiments(Baskin et al., 1994). (**F**) Scale bars of auxin concentration and cell growth rate.

Next, we tested how local mechanical disruptions of QC, root tip and LRC would alter the model outcome, and whether this outcome agrees with experimental observations. The QC is a small group of cells (four to seven in the *A. thaliana* root) located in the middle of the root apical meristem(Doerner, 1998). The QC divides infrequently and grows at an extremely low rate(Nawy et al., 2005). The QC is known to be the source of signals that inhibits differentiation of the surrounding stem cells(Van Den Berg et al., 1997). QC cells define the correct location of the stem cell niche but also behave as independent cells by self-renewing and replenish initials that have been displaced from their position(Kidner et al., 2000). QC laser ablation is not lethal for the root as a new QC and stem niche is quickly reconstructed a few cells above the previous one as a consequence of auxin accumulation(Sabatini et al., 2003). We replicated the same experiment *in silico* by removing the two QC cells from the model after a few hours of simulation (Supplementary Fig. S12C and Supplementary Video S9). Compared to the simulated wild-type (Fig. 5A), the typical auxin accumulation in the root tip is depleted, and auxin reflux in the LRC is significantly reduced, while most of auxin coming from the shoot tends to concentrate in the cells above the lost QC, exactly as observed in experiments (see Fig. 5 in (Reddy et al., 2007)). Similarly, removal of LRC led to sharp auxin accumulation in the root tip (Fig. 5C and Supplementary Video S10), largely matching empirical reports(Tsugeki and Fedoroff, 1999).

*A. thaliana* roots are able to survive not only after QC ablation but even after the complete excision of the root tip, as the plant is able to regenerate a new root tip including a complete new root apical meristem(Efroni et al., 2016). Since the stem cell niche is lost with excision, the regeneration process relies on other pluri-potent dormant cell types available in the remaining stump(Sugimoto et al., 2010). We tested if we could replicate this experiment by removing the entirety of the root tip during the simulation (Fig. 5D and Supplementary Video S11). Compared to QC ablation (Supplementary Fig. S12C and Supplementary Video S9), the removal of the root tip displays an even more radical effect on the auxin patterning dynamics (Fig. 5D and Supplementary Video S11). Auxin signal was strongly increased in the vascular tissues and auxin reflux in the lateral tissues was absent; again, model predictions closely match experimentally observed patterns (see Fig. 1 in (Matosevich et al., 2020)).

Finally, we explored whether simulated chemical perturbation of cytoskeleton would predict experimentally observed root phenotype. CMTs organization can be modulated by chemical treatments which cause microtubules depolymerization and stimulate the radial expansion of roots(Baskin et al., 1994). We simulated CMTs disruption by implementing a gradual degradation of CMTs during root growth (Fig. 5E and Supplementary Video S12). The simulated root growth displays a marked radial swelling, more evident in the center of the meristem and much less pronounced in the root tip (Fig. 5E). Several cells divide irregularly, as a result of loss of anisotropic growth. As a consequence, PINs localizations on the membrane became irregular and asymmetric auxin distribution was eventually lost (Fig. 5E).

These results demonstrate that our framework can be used to predict various root meristem phenotypes including several auxin transport mutants, and mechanical or chemical manipulations of root tissue geometry and thus it is a powerful and rapid resource for guiding new experimental designs in the laboratory.

## Discussion

Today, computer models have become a central tool for wet-lab scientists to quickly explore possible mechanisms underlying morphogenesis before designing and implementing effective experimental strategies. To date, computer models of root development have been instrumental in understanding root maturation in the context of biochemical flow over non-growing(Band et al., 2014; Rutten and ten Tusscher, 2019) or idealized templates(Grieneisen et al., 2007). On contrary, little to no attempts were made to couple mechanisms of cell polarity establishment and realistic tissue biomechanics at single cell resolution to mechanistically understand how a root emerges from small populations of cells.

Here, we took advantage of an efficient technique called the Position-Based Dynamics(Müller et al., 2007) to resolve biomechanics of root growth, while simultaneously incorporating detailed biochemical reactions that guide auxin production, distribution and transport at the subcellular level. Our mechanistic cell-based framework successfully predicted the root meristem morphology including tissue patterns, polarity organization, auxin distribution and anisotropic growth from the embryonic onset. Tissue-level properties can emerge from local cell activities and direct cell-to-cell communication through mobile auxin signal with no need for global regulators or polarizers. Anisotropic cell growth and PINs localizations self-organize in complex structures such as the root tip and the cortex-epidermis interface. Furthermore, our model demonstrates that auxin influx from the LRC and subsequent ‘bipolar’ PINs localization in the cortex tissues may be important elements of the auxin reflux-loop system that is robust under mechanical and growth constrains. This system can sustain root growth over prolonged time periods, whereas root growth anisotropy would emerge naturally from differential growth rate of adjacent tissues.

Finally, we propose that the complexity of PIN polarities could result from dynamic cooperation between auxin flow and biomechanical cues, and we suggest a plausible molecular mechanism for cell polarization based on a putative kinase/phosphatase regulation(Hajný et al., 2020; Michniewicz et al., 2007; Weller et al., 2017). The current model can be extended to address many aspects of root development and organogenesis including root gravitropism, root-tip regeneration after ablation, root response after chemical treatment and mutant simulations. In the future, our model can be expanded to address additional mechanisms of root zonation(Ivanov and Dubrovsky, 2013), stem cells differentiation(Sabatini et al., 2003), lateral root initiation(Perianez-Rodriguez et al., 2021), and auxin flux through plasmodesmata(Mellor et al., 2020), indicating the possibility for modular extensions of the current framework to account for further complexities; for instance the action of other hormones and postembryonic regulatory mechanisms like gravitropism and phototropism.

As the quantitative model predictions are in a fair agreement with experimental observations, our framework will be very useful to predict the phenotypes of various root mutants and test the effects of perturbations such as chemical treatments, gene knockdown or mechanical alterations, thereby fostering the effective design of experiments in the laboratory. This work combines different aspects of morphodynamics and represents a leap forward towards quantitative models of plant organogenesis which could serve as a next-stage platform to develop novel traits of high socio-economic importance.

## Materials and Methods

### *Arabidopsis thaliana* embryo cell segmentation

In order to obtained the segmented digital mesh of an embryo template it is necessary to observe the following procedure: 1) Get the microscope picture of an *A. thaliana* embryo without the aerial parts, segmentation is better obtained from pictures with a black background and colour cell borders with high contrast; 2) Load the picture in the MorphoGraphX Stack; 2) Go to the view tab and turn on the cutting surface. Click the grid checkbox and hit reset. It should draw a plan around the image. You can use the sliders to adjust the size; 3) Use the Mesh-Creation-Mesh Cutting Surface process. This will create a mesh in that location. Turn off the cutting surface. 4) Subdivide the mesh a few times; 5) Project signal, using a range like (−1, 1) as the mesh should be in the middle. If the projection is too coarse, subdivide the mesh again; 6) Segment the stack as described in the MorphoGraphX tutorial(de Reuille et al., 2015); 7) Convert the mesh into a cell mesh with Tools-Cell Maker-Mesh 2D-Tools-Polygonize Triangles. Set the “Max Length” parameter to zero; 8) a final pass of smoothing can help to remove the jagged lines from the cells; 9) rescale the mesh to the desired size and save it.

### Computational model

The model was created using MorphoDynamX, the next generation of the MorphoGraphX software(de Reuille et al., 2015). This modelling framework is based on an advance data structure called Cell Complexes(Karwowski and Prusinkiewicz, 2004; Prusinkiewicz and Lane, 2013) that overcomes the typical limitations of previous technologies like Vertex-Vertex complexes(Federl and Prusinkiewicz, 1999) and are ideal for modelling subdividing geometry in 2 and 3 dimensions. MorphoDynamX provides the user interface and application program interface to the cell complex structures, on top of which the current model has been developed. Cells are represented as triangulated polygons obtained through the segmentation process described in the previous section. Cells are composed of vertices connected by edges, and bounding internal faces. Each of these three elements (vertices, edges and faces) has their own biological interpretation and possesses different attributes and properties that allow the model to run and produce a root-like structure. Perimeter edges represent the cell walls while internal edges might be broadly identified as the cell cytoskeleton. The model does not distinguish between cell walls and cell membranes, both are depicted as the same edge but they are computationally distinguished depending on the situation (whether calculating mechanical or chemical effects). Cells at the very top of the model are considered either sources or sinks (in the real heart stage of the embryo auxin flows into the epidermal layer of the upper part and again back into the vascular tissues of the root, creating a auxin loop inside the embryo), so auxin that reaches the top is automatically removed from the system. The mechanical growth of the simulated root is based on Position-Based Dynamics (PBD)(Müller et al., 2007). Position-based dynamics is a method used to simulate physical phenomena like cloth, deformation, fluids, fractures, rigidness and much more(Müller et al., 2007). PBD allows overcoming the typical limitations of force-based models by omitting the velocity layer and immediately working on the positions following constraints that restrict the systems dynamics in order to obtain a desired effect. Chemical processes are numerically solved using the simple Euler method (Butcher, 2007). Model implementation is described in details in the Supplementary Information section.

### Time-lapse confocal imaging of young seedlings

Confocal laser-scanning micrographs of 35S::PIP2-GFP transgenic lines were obtained as published elsewhere^82^. Briefly, seeds were stratified for 3 days, seed coat was removed and peeled embryos were imaged using vertical Zeiss LSM700 microscope with a 488-nm argon laser line for excitation of GFP fluorescence. Emissions were detected between 505 and 580 nm with the pinhole at 1 Airy unit, using a 20x air objective. Images were taken every 20 minutes and Z-stack maximal projections were done using ImageJ software.

## Acknowledgments

We are grateful Richard Smith, Anne-Lise Routier, Crisanto Gutierrez and Juergen Kleine-Vehn for providing critical comments on the manuscript. Funding: This work was supported by the Programa de Atraccion de Talento 2017 (Comunidad de Madrid, 2017-T1/BIO-5654 to K.W.), Severo Ochoa (SO) Programme for Centres of Excellence in R&D from the Agencia Estatal de Investigacion of Spain (grant SEV-2016-0672 (2017-2021) to K.W. via the CBGP). In the frame of SEV-2016-0672 funding M.M. is supported with a postdoctoral contract. K.W. was supported by Programa Estatal de Generacion del Conocimiento y Fortalecimiento Cientıfico y Tecnologico del Sistema de I+D+I 2019 (PGC2018-093387-A-I00) from MICIU (to K.W.). MG is recipient of an IST Interdisciplinary Project (IC1022IPC03).

