## Supplementary Information for "A coupled mechano-biochemical framework for root meristem morphogenesis"

### Supplemental Information

#### 1 General Model Assumptions

The root of *A. thaliana* is made of several radially-organized layers of morphologically similar cells that can be easily distinguished by visual inspection of both radial and cross sections(Dolan et al., 1993; Scheres et al., 1994). The central vascular tissue is composed by a bundle of thin and elongated cells surrounded by the pericycle, a cylindrical sheath protecting the stele. The pericycle is also the origin of emerging lateral organs(Lavenus et al., 2013; Péret et al., 2009). The central cylinder (stele) is enclosed by three adjacent tissues endodermis, cortex and epidermis. The gravity-sensing columella is located at the very tip of the root and it is composed of four layers of differentiated cells(Kumpf and Nowack, 2015). The meristem of the mature root is covered by the lateral root cap which protects the meristem and is periodically shed and replaced by new emerging layers(Di Mambro et al., 2019; Kumar and Iyer-Pascuzzi, 2020). Finally, the root tip stores a group composed of undifferentiated stem cells that divide asymmetrically and replenish the upper sections of individual tissues(Stahl and Simon, 2009). Therefore, this precise spatio-temporal arrangement of tissues in the root requires the coordination of cell polarity, anisotropic growth and asymmetric cell divisions. However, mechanisms underlying this coordination are poorly understood.

Auxin-driven root growth of *A. thaliana* has been intensively studied in the last years, and it is known to be one of the major players in root development(Ljung, 2013). Auxin distributes along the root through a tightly controlled mechanism and its disruption results in the organ growth failure(Truman et al., 2010). Auxin synthesis and homeostasis is thought to be the other major contributor to cell elongation(Velasquez et al., 2016). The main source of auxin during globular root embryogenesis comes from the shoot, and tends to accumulate in vascular tissue, root tip and epidermis(Robert et al., 2015; Smit and Weijers, 2015). Many aspects of auxin transport by PIN efflux carriers are well understood, however the connections between these mechanisms and the effect on root growth are largely obscure(Adamowski and Friml, 2015; Habets and Offringa, 2014).

Given the previously described biological properties of *A. thaliana* root, we decided to integrate the following biological observations:

1. The root is composed of cells categorized into different lineages: QC, Columella Initial, Columella, Epidermal/LRC initial, Cortex/Endodermis Initial (CEI), Cortex/Endodermis Initial daughter (CEID), Lateral Root Cap (LRC), Epidermis, Endodermis, Cortex, Pericycle and Vascular. The latter includes all central vascular tissues, for simplicity. Cell types are not uniform and each of them can possess different properties and display specific behaviors(Benfey et al., 2010; Nawy et al., 2005). For example, each cell type expresses preferentially different PIN protein families(Paponov et al., 2005). These characteristics are discussed in more detail later.
2. Cells are represented as triangulated polygons. Perimeter edges represent the cell walls while internal edges can be broadly identified as the cell cytoskeleton and internal structure. Currently, there is no distinction between cell membranes and cell walls, both are represented by a single edge, which are referred to as "membrane sections" in the text.
3. Cell expansion is described according to the acid-growth hypothesis(Rayle and Cleland, 1992). Cells are under constant osmotic pressure, and their expansion is prevented by a stiff cell wall with viscoelastic properties. Cells can be considered as incompressible objects. Cell walls possess a strong extensional stiffness at very low or negligible auxin concentration (but also at very high auxin concentration, as discussed later), which prevents the cell from expanding. Auxin (indole-3-acetic acid, IAA), induces acidification of the cell wall activating a range of enzymatic reactions which modifies the extensibility of plant cell walls, allowing the cell to grow(Cosgrove, 2000; Hager et al., 1991).
4. Mechanical strain induces cortical microtubule (CMTs) orientation and consequently restricts growth, orthogonal to CMTs. It has been shown that mechanical stress can affect microtubules disposition inside the cell(Hamant et al., 2019). Microtubules restrict cell expansion in a determined direction allowing anisotropic growth(Daher et al., 2018).
5. Cell division occurs according to cell polarity, cell type and condition.
6. Auxin flows into the root from the aerial section of the plant through the vascular tissues (Petrášek and Friml, 2009). Auxin can also be locally produced in the root apical meristem(Kerk et al., 2000).
7. Auxin is passively diffused inside cells and all over the intercellular space. Auxin is also actively transported by auxin carriers(Hošek et al., 2012). Auxin exchange between cells is not direct but it occurs through the intercellular space, which is not visually displayed but still considered during modelling computations.
8. Auxin induces PINs and AUX1/LAX protein expression(Zwiewka et al., 2019). PINs are subsequently trafficked to the cell membranes according to several criteria as discussed later. Briefly, CMTs and auxin flux/concentration are the main contributors to PINs localization.

The previous assumptions and their implications are therefore merged into a comprehensive and coherent model structure (Supplementary Fig. S4 and Supplementary Fig. S5). Details about each model part are explained in the following sections. Optimal parameters values were chosen after testing over a large plausible range of values for each parameter. Parameters description and value are listed in Supplementary Table S1.

#### 2 Cell Polarization Model

The processes that define cell polarity in plants are not well understood(Dettmer and Friml, 2011), and are considered to be different than those in animals. Plant cells display clear polarity patterns when observed to grow anisotropically or by targeting proteins to certain parts of the cell membranes(Yang, 2008). Root cells present a clear apical-basal (shootward-rootward) polarity, which allows them to elongate towards a determined direction pumping hormones and other molecules creating desired flows(Kleine-Vehn and Friml, 2008). Almost all cell types in *A. thaliana* root display a clear anisotropic pattern(Baskin, 2005). Cell polarity also strongly correlates with PINs localization(Willemsen and Keren, 2003). Up to the date, several prominent polarity markers have been identified in plants, among those are PIN auxin efflux carriers(Wisniewska et al., 2006), and putative regulators of cell division orientation BASL(Pillitteri et al., 2011) and SOSEKI(Yoshida et al., 2019). Regarding PIN proteins, the only known regulators that directly impact on their polarity are the AGCVIII kinases and PP2A phosphatases(Barbosa et al., 2018), components of the phosphorylation on/off switch aimed at the central hydrophilic loop of PINs(Michniewicz et al., 2007). Auxin transport, mechanical stress, nutrients availability and pathogen response are other processes known to contribute to the cell polarity modulation(Adamowski and Friml, 2015). It has been recently shown that external stress can affect internal microtubules organization and therefore guide the cell towards a determined polarity(Hamant et al., 2019). During root swelling the cells are growing isotropically but also undergo stretching in a direction determined by their geometry and their position inside the organ. According to previous observations, cell microtubules will align perpendicular with the maximal strain, hence enforcing polarity(Hamant et al., 2008). Once polarity has been clearly determined the cell will acquire anisotropic growth and will enlarge only in a specific direction.

We integrated these findings into a comprehensive polarity system assuming the following:

1. Cortical microtubules orientation restricts the growth anisotropy, orthogonal to the CTMs.
2. External mechanical stresses affect the orientation of microtubules, therefore inducing changes in growth orientation. Microtubules arrange orthogonally to wall strains. In a sense it can be said that the cell “follows” the imposed stress by regulating its own polarity according to the external stresses, using walls strains as reference. Microtubules orientation is represented as a single vector for each cell and, as a consequence, cell polarity can be easily defined as the orthogonal vector to the latter.
3. Cortical microtubules reorientation is triggered only if a certain strain threshold is reached, therefore simulating a plastic behavior.

4. For sake of simplicity, we assume that after cell division newborn cells initially inherit the same CMTs orientation as the mother.
5. A dynamic microtubule reorganization is defined by the following formula:

$$\frac{d\,\overrightarrow{MF_{cell}}}{dt} = \overrightarrow{MF_{cell}} + R_{MF} \sum_{i}^{m} u\left(\overrightarrow{MF_{cell}}\right) abs\left(\left(\overrightarrow{u(MF_{cell})} \cdot \overrightarrow{u(mem_i)}\right) \epsilon_{mem_i}\right) - d_{MF}\,\overrightarrow{MF_{cell}} \qquad (1)$$

(1) $\overrightarrow{MF_{cell}}$ is the vector denoting microtubules orientation inside the cell; $R_{MF}$ is microtubules reorientation rate; *m* is the total number of membrane sections; $u(\overrightarrow{MF_{cell}})$ is the unit vector parallel to the microtubules vector; $u(mem_i)$ is the unit vector parallel to membrane section $mem_i$ ; $\epsilon_{mem_i}$ is the strain rate of membrane section $mem_i$ ; $d_{MF}$ is microtubules decay rate. The dot "·" symbol indicates the dot product between vectors.

#### 2.1 Auxin Transport Model

As already discussed in the main text, previously proposed models of auxin polar transport can be approximately divided into two main strategies: flux-based and concentration-based methods(van Berkel et al., 2012). Broadly speaking, flux-based models assume that PIN polarize according to the direction of auxin influx into the cell in order to reinforce the flow in that same direction(Alim and Frey, 2010; Feugier et al., 2005; Feugier and Iwasa, 2006; Fujita and Mochizuki, 2006; Mitchison, 1980; Stoma et al., 2008). In concentration-based models it is assumed that the cell is able to detect auxin concentration levels and to trafficking PIN either against or along the gradient(Jönsson et al., 2006; Merks et al., 2007; Newell et al., 2008; Smith et al., 2006). A transport feedback model closely related to the flux-based model for leaf venation patterning was proposed a few years ago(Kramer, 2009; Smith and Bayer, 2009). Despite relying on different formulations, both model types are able to recreate many auxin-related patterns observed in nature. Phyllotaxis, which creates regularly spaced spots around the shoot apical meristem, has been modelled both by diffusing inhibitors(Hellendoorn and Lindenmayer, 1974) and reaction–diffusion models(Meinhardt, 2008). However, the models that appear to best represent available data about auxin and its transporters point to a transport-feedback mechanism(De Reuille et al., 2006; Heisler et al., 2005; Jönsson et al., 2006; Smith et al., 2006; Smith and Bayer, 2009). Phyllotaxis and leaf venation patterns appear very different, and require mechanisms with different properties. Recent years have witnessed the emergence of appealing alternative theoretical hypotheses for polar auxin transport. One of these is the extracellular receptor-based polarization (ERP) model(Wabnik et al., 2010). In this framework, PINs polarity could be regulated by auxin receptors in the apoplast that in turn inhibit local PINs endocytosis after binding auxin. The ERP model easily reproduces several processes like de novo vascularization, venation patterning, and tissue regeneration with only minimal initial assumptions. A mechanistic alternative model was proposed by Heisler(Heisler et al., 2010). The author noticed a clear correlation between PINs polarity and the alignment of cortical microtubules, which led him to assume that wall stress induced by auxin accumulation could be involved in determining PINs localization. A more recent study(Narasimhan et al., 2021) showed that auxin has a positive, PIN2-specific effect on PIN2 endocytosis, implying that auxin may have a negative feedback on PIN2 localization.

In the current study we focused on flux-based and concentration-based model alternatives and verified their effect on auxin transport regulation. In our model both PINs and AUX1/LAX expression is induced by the presence of auxin inside the cell. PIN trafficking to the membranes is also regulated by auxin, while it is kept constant for AUX1. Auxin is exported by PINs from the cell into an intercellular space (not visualized in the simulation), from which it can be imported by AUX1 from neighbouring cells. This allows adjacent cells to continuously exchange auxin. Given the amount of available evidence, we incorporated following assumptions regarding auxin transport in roots:

1. As already discussed, the cell membrane is represented by a two-dimensional polygon. Each edge of the polygon denotes a section of the cell wall/membrane (indicated as $mem$ in the following formulas). Each membrane section possesses its own mechanical and chemical attributes, such as PIN and AUX1/LAX amounts, mechanical strain and orientation. Each membrane section possesses an attributes indicating the intercellular auxin contained between two neighboring cells.
   Auxin and protein species are represented in the following formulas as concentrations, that is, number of molecules divided by the area of the cell or the intercellular space. For example $IAA_{cell} = molecules\ of\ IAA/area\ of\ the\ cell$
2. Auxin is imported by AUX/LAX from the intercellular space and exported in a polar manner from cells by PINs with the cooperation with PGP1/ABC transporter family(Geisler and Murphy, 2006). In the current model we do not consider the PGP1/ABC transporters, and active auxin transport is regulated only by PIN and AUX/LAX carriers:

$$I_{mem} = K_{AUX1}\ AUX1_{mem}\ IAA_{mem}\ L_{mem} \quad (2)$$

$$E_{mem} = K_{PIN}\ PIN_{mem}\ IAA_{cell}\ L_{mem} \quad (3)$$

$$\frac{dIAA_{mem}}{dt} = \sum_{i}^{n}(E_{mem} - I_{mem})_{cell_i} - \sum_{i}^{m} DI_{IAA}\frac{IAA_{mem} - IAA_{mem_i}}{L_{mem} + L_{mem_i}} - D_{IAA}(IAA_{cell}) - d_{IAA}(IAA_{mem}) \quad (4)$$

$$\frac{dIAA_{cell}}{dt} = b_{IAA} + \sum_{i}^{m}(I_{mem} - E_{mem})_{cell_i} + D_{IAA}(IAA_{cell}) - d_{IAA}(IAA_{cell}) \quad (5)$$

(2) $I_{mem}$ is the auxin imported into the cell through a specific membrane section *mem*; $K_{AUX1}$ is the coefficient of auxin importing rate by AUX1/LAX; $AUX1_{mem}$ is the amount of AUX1/LAX protein localized on membrane section *mem*, $IAA_{mem}$ is the intercellular auxin available in the membrane section *mem*.

(3) $E_{mem}$ is the auxin exported from the cell into the intercellular space through a specific membrane section *mem*; $K_{PIN}$ is the coefficient of auxin exporting rate by PIN; $PIN_{mem}$ is the amount of PIN protein localized on membrane section *mem*, $IAA_{cell}$ is the concentration of auxin inside the cell; $L_{mem}$ is the length of membrane section *mem*.

(4) $IAA_{mem}$ is the intercellular auxin available by the current membrane section *mem;* $IAA_{mem_i}$ is the amount of intercellular auxin available by a neighbor of the current membrane section *mem;* $cell_i$ is a cell sharing the current membrane section *mem*; $mem_i$ is a membrane section neighbouring the current membrane section *mem;* $DI_{IAA}$ is the auxin diffusion rate in the intercellular space; $L_{mem}$ is length of membrane section *mem;* $L_{mem_i}$ is length of the neighbouring membrane section *mem*$_i$*;* $d_{IAA}$ is the auxin degradation for the intercellular auxin available by the current membrane section *mem* (described in section 4).

(5) $IAA_{cell}$ is the auxin concentration inside the current *cell;* $b_{IAA}$ is the auxin basal production rate; $D_{IAA}(IAA_{cell})$ is the net auxin diffusion for the current *cell* (this functions is described in section 3); $d_{IAA}(IAA_{cell})$ is the auxin degradation for the current *cell* (this function is described in section 4).

3. Auxin can diffuse passively into cells from the intercellular space(Petrášek and Friml, 2009) according to the formula:

$$D_{IAA}(IAA_{cell}) = \sum_{i}^{m} P_{IAA}(IAA_{mem_i} - IAA_{cell})\, L_{mem_i} \qquad (6)$$

(6) $D_{IAA}(IAA_{cell})$ is the net auxin diffusion between the cell and the surrounding intercellular space; *m* is the total number of membrane sections; $P_{IAA}$ is membrane permeability; $IAA_{mem_i}$ is the intercellular auxin available by membrane section *mem*$_i$ ; $IAA_{cell}$ is the auxin concentration inside the cell; $L_{mem_i}$ is the length of membrane section *mem*$_i$.

4. Auxin decays following the combined effect of conjugation and oxidation, at a constant rate(Ljung, 2013). Above a determined concentration threshold of auxin inside the cell or a membrane section *mem*, auxin degradation is increased:

$$d_{IAA}\left(IAA_{cell/mem}\right) = d_{IAAb} + (d_{IAAMax} - d_{IAAb}) \frac{{IAA_{cell/mem}}^4}{{K_{IAAMax}}^4 + {IAA_{cell/mem}}^4} \qquad (7)$$

(7) $d_{IAA}\left(IAA_{cell/mem}\right)$ is the auxin degraded inside a cell or in the intercellular space available to section *mem*; $d_{IAAb}$ is the basal auxin degradation rate; $d_{IAAMax}$ is the maximum auxin degradation rate; $IAA_{cell/mem}$ is the current auxin concentration inside the cell or available to a membrane section *mem*; $K_{IAAMax}$ is the coefficient for half-max auxin degradation.

5. Auxin regulates the expression of the auxin carriers(Heisler et al., 2005), by up-regulating PINs and AUX1/LAX expressions:

$$\frac{dAUX1_{cell}}{dt} = b_{AUX1} + AUX1_{Expr} \frac{{IAA_{cell}}^2}{{AUX1_K}^2 + {IAA_{cell}}^2} - AUX1_{cell}\, AUX1_{tr} - d_{AUX1} AUX1_{cell} \qquad (8)$$

$$\frac{dPIN_{cell}}{dt} = b_{PIN} + PIN_{Expr}\frac{{IAA_{cell}}^2}{{PIN_K}^2 + {IAA_{cell}}^2} - PIN_{cell}\,PIN_{tr} - d_{PIN}PIN_{cell} \quad (9)$$

(8) $AUX1_{cell}$ is the cytoplasmic AUX1/LAX inside the cell; $b_{AUX1}$ is AUX1/LAX basal expression; $AUX1_{Expr}$ is the auxin-induced AUX1/LAX maximal expression; $AUX1_K$ is the auxin-induced AUX1/LAX half-max expression; $IAA_{cell}$ is the auxin concentration inside the cell; $d_{AUX1}$ is AUX1/LAX degradation rate. AUX1/LAX expression is disabled when the maximum concentration $AUX1_{Max}$ (see Table S1) is reached in the cell.

(9) $PIN_{cell}$ is the cytoplasmic PINs inside the cell; $b_{PIN}$ is the PIN basal expression; $PIN_{expr}$ is the auxin-induced PIN maximal expression; $PIN_K$ is the auxin-induced PIN half-max expression; $IAA_{cell}$ is the auxin concentration inside the cell; $d_{PIN}$ is PIN degradation rate. PIN expression is disabled when the maximum concentration $PIN_{Max}$ (see Table S1) is reached in the cell.

6. Auxin modulates sub-cellular localization of its exporters(Sauer et al., 2006). AUX1/LAX is redistributed evenly among the membrane sections making part of the cell membrane, while PINs are redistributed depending on the "PIN sensitivity" of each membrane section (as described in section 7):

$$\frac{dAUX1_{mem}}{dt} = AUX1_{cell}\,AUX1_{tr}\frac{L_{mem}}{\sum_i^m L_{mem_i}} - d_{AUX1}AUX1_{mem} \quad (10)$$

$$\frac{dPIN_{mem}}{dt} = PIN_{cell}\,PIN_{tr}\,PinS_{mem} - d_{PIN_{mem}}PIN_{mem} \quad (11)$$

$$d_{PIN_{mem}} = d_{PIN} + (d_{PINmax} - d_{PIN})\frac{1}{1 + IAA_{cell}} \quad (12)$$

(10) $AUX1_{mem}$ are the AUX1/LAX proteins on membrane section *mem*; $AUX1_{cell}$ is the concentration of AUX1/LAX in the cytoplasm; $AUX1_{tr}$ is AUX1/LAX trafficking rate; *m* is the total number of membrane sections; $L_{mem}$ is the length of membrane section *mem*; the element $L_{mem}/\sum_i^m L_{mem_i}$ therefore indicates the fraction of cytoplasmic AUX1/LAX trafficked to the membrane section *mem*; $d_{AUX1}$ is AUX1/LAX degradation rate. AUX1/LAX trafficking is disabled when the maximum concentration permitted on one unit of membrane section $AUX1_{Maxmem}$ (see Table S1), is reached.

(11) $PIN_{mem}$ are the PIN proteins on membrane section *mem*; $PIN_{cell}$ is the concentration of PIN in the cytoplasm; $PIN_{tr}$ is PIN trafficking rate; $PinS_{mem}$is the current PINs sensitivity of membrane section *mem,* which determines the fraction of cytoplasmic PINs that are trafficked to membrane section *mem* (see section 7 later for a detailed description of this term); $d_{PIN_{mem}}$ is PIN degradation dynamic formula on the membranes, described in equation 12. PIN trafficking is disabled when the

maximum concentration permitted on one unit of membrane section $PIN_{Maxmem}$ (see Table S1), is reached.

(12) $d_{PINmem}$ is PIN degradation dynamic formula on the membranes; $d_{PIN}$ is PIN base degradation rate (same as cytoplasmic degradation); $d_{PINmax}$ is the maximum PIN degradation on the membranes; $IAA_{cell}$ is the current auxin concentration inside the cell; PINs degradation on the membranes is modelled in this dynamic way in order to allow for a faster turnout of the protein from the membranes when auxin level in the cell is low(Kleine-Vehn et al., 2008).

7. PINs are trafficked to the membranes according to a specific criterion; each membrane section *mem*, possesses an instrumental "PIN sensitivity" property, which regulates the propensity of that membrane section to incorporate additional PINs. This property is a phenomenological parameter for more low-level processes involved in PINs trafficking, like PINs phosphorylation and endocytosis. PIN sensitivity is always a number between 0 and 1 and the PIN sensitivity from all membranes sections sum up to 1, representing the total amount of cytoplasmic PINs that it is trafficked to the membranes. PIN sensitivity is the linear combination of auxin flow (defined either by auxin flux or auxin concentrations), growth anisotropy (defined by CMTs orientation), and cell geometry. Cell geometry in fact is known to be an important regulating factor of protein trafficking(Elliott and Kirchhelle, 2020). Columella cells are the only exception to this rule; in the columella PINs are always trafficked uniformly among membrane sections, in order to reflect the observed PIN3 distribution(Friml et al., 2002). PIN sensitivity for a given membrane section is defined as:

$$PinS_{mem} = \frac{\exp\left(PinSR_{mem}\right)}{\sum_i^m \exp\left(PinSR_{mem_i}\right)} \quad (13)$$

$$PinSR_{mem} = kMF\ IMF_{mem} + kP\ IP_{mem} + kMFP\ (IMF_{mem} + IP_{mem}) + kG\ IG_{mem} \quad (14)$$

$$IMF_{mem} = \left|\overrightarrow{MF_{cell}}\right| \frac{U\left(\overrightarrow{MF_{mem}}\right)^4}{U\left(\overrightarrow{MF_{mem}}\right)^4 + Kmf^4};\ U\left(\overrightarrow{MF_{mem}}\right) = \overrightarrow{u(MF_{cell})} \cdot \overrightarrow{n(mem)} \quad (15)$$

$$IG_{mem} = \left(\overrightarrow{u(axisMax)} \cdot \overrightarrow{n(mem)}\right) \frac{\left(1-\frac{\left|\overrightarrow{axisMin}\right|}{\left|\overrightarrow{axisMax}\right|}\right)^4}{\left(1-\frac{\left|\overrightarrow{axisMin}\right|}{\left|\overrightarrow{axisMax}\right|}\right)^4 + Kgeom^4} \quad (16)$$

(13) PIN sensitivity for membrane section *mem* is obtained by applying the soft-max function over all the raw PIN sensitivities $PinSR_{mem_i}$ calculated for each membrane section of the cell. The soft-max function was used to normalize the total sum of raw sensitivities to 1.

(14) We define the raw PIN sensitivity for a membrane section *mem*. $IMF_{mem}$ is the contribute of cortical microtubules orientation to PIN sensitivity for membranes section *mem* (see the equation 15); $kMF$ is the coefficient of contribution of CMTs orientation to PIN sensitivity; $IP_{mem}$ is the contribute of auxin flow to PIN sensitivity

for membrane section *mem* (this parameter depends on the chosen method between auxin-flux and auxin-concentration, see the following section); $kP$ is the coefficient of contribution of auxin flow to PIN sensitivity; $kMFP$ is the coefficient of interaction of the two contribution from CMTs orientation and auxin flow to PIN sensitivity; $IG_{mem}$ is the contribution of cell geometry to PIN sensitivity for membranes section *mem* (*s*ee the equation 16); $kG$ is the coefficient of contribution of cell geometry to PIN sensitivity.

(15) We define the impact of CMTs orientation $IMF_{mem}$ on PIN sensitivity of membranes section *mem*. $\overrightarrow{MF_{cell}}$ is the vector describing CMTs orientation in the cell (defined in equation 1); $U(\overrightarrow{MF_{mem}})$ is the normalized effect of CMTs orientation on membrane section *mem*; $\overrightarrow{u(MF_{cell})}$ is the unit vector parallel to the microtubules vector; $\overrightarrow{n(mem)}$ is the unit vector orthogonal to membrane section *mem*, this is an intermediate value to calculate $IMF_{mem}$; $Kmf$ half-max contribute of CMTs orientation to PIN sensitivity. The dot "·" symbol indicates the dot product between vectors.

(16) We define the impact of cell geometry on PIN sensitivity of membranes section *mem*; $\overrightarrow{n(mem)}$ is the unit vector orthogonal to membrane section *mem*; $\overrightarrow{axisMax}$ and $\overrightarrow{axisMin}$ are the longest and the shortest cell principal axis, respectively; $\overrightarrow{u(axisMax)}$ is the unit vector parallel to the longest cell axis; $Kgeom$ is the half-max cell geometry contribution to PIN sensitivity. The dot "·" symbol indicates the dot product between vectors.

As discussed before, PIN sensitivity of a specific membrane section is affected either by auxin flux or auxin concentration, depending on the "auxin-flux' or 'regulator-polarizer' scenario. The chosen model will determine the $IP_{mem}$ reaction introduced in the previous section in equation (14).

##### 2.1.1 Auxin-flux method

Flux-based models of auxin transport were first introduced by Mitchison(Mitchison, 1980), which stated that cells sensing the flux of a molecule in a certain direction increases their capacity to transport the molecule in that same direction. This type of strategy is well suited to reproduce situations where auxin displays a strong canalized patterning like in vein formation(Rolland-Lagan and Prusinkiewicz, 2005). Flux-based models assume the existence of cellular flux-sensing mechanisms that have not been discovered so far, and therefore the biological plausibility of these models is a matter of debate. In the auxin-flux model, we calculate the net vector of auxin flow and redirect PINs accordingly. Specifically, the total auxin-flux vector for a cell is defined as:

$$\overrightarrow{FLUX_{cell}} = \sum_{i}^{m} \overrightarrow{u(centroid_{cell}, midpoint_{mem_i})}\,(I_{mem_i} - E_{mem_i}) \qquad (16)$$

(16) *m* is the total number of membrane sections; $centroid_{cell}$ is the centroid of the cell, $midpoint_{mem_i}$ is the midpoint of membrane section *mem*; $\overrightarrow{u(centroid_{cell}, midpoint_{mem_i})}$ is the

unit vector parallel to the vector connecting the two previous elements; $\left(I_{mem_i} - E_{mem_i}\right)$ indicates the net amount of auxin crossing the membrane section *mem*.
Given the auxin-flux vector for a cell, the PIN sensitivity contribution by auxin flow on a given membrane section *mem* as required in equation (14) can be obtained by projecting the cell flux over the section *mem* in order to get the section-specific flux, and next by calculating the actual contribution:

$$IP_{mem} = \frac{{F_{mem}}^4}{{F_{mem}}^4 + Kflux^4};\ F_{mem} = \left(\overrightarrow{FLUX_{cell}} \cdot \overrightarrow{n(mem)}\right) L_{mem} \qquad (17)$$

(17) $IP_{mem}$ is the (unit less) auxin flow contribution to PIN sensitivity (for the auxin-flux method); $F_{mem}$ is the effect of the cellular auxin-flux vector to membrane section *mem*; $Kflux$ is the auxin-flux effect half-max contribution on PIN sensitivity; $\overrightarrow{FLUX_{cell}}$ is the cellular auxin-flux vector; $\overrightarrow{n(mem)}$ is the unit vector orthogonal to the membranes section *mem* and $L_{mem}$ is the length of the membrane section *mem*. The dot "·" symbol indicates the dot product between vectors.

##### 2.1.2 Regulator-Polarizer method

In a concentration-based model the cell is able to sense the difference in auxin concentration between itself and the neighboring cells and allocate PINs on the membrane accordingly(van Berkel et al., 2012). This kind of model assumes the existence of short-distance signaling between cells and is usually designed to work up-the-gradient. Given the fact that such signaling molecules have not been so far discovered, an alternative model was proposed(Wabnik et al., 2010) where auxin imported through the cell wall binds to a specific receptor and inhibits endocytosis of PINs from the nearest membrane. This adaptation makes the model more flexible and able to reproduce down-the-gradient auxin flow like in flux-based models, as well as being more biologically sound than auxin-flux models described before.
Starting from this intuition, we developed a concentration-based mechanism for auxin transport capable of reproducing a self-organizing polarization. Cell polarization have been investigated both on theoretical ground(Gierer and Meinhardt, 1972; Jilkine and Edelstein-Keshet, 2011; Meinhardt and Gierer, 2000) and designed synthetic circuits(Chau et al., 2012; Rappel and Edelstein-Keshet, 2017). In our framework, PIN polarity emerges from the interaction between three molecules: auxin, a polarizer and a regulator. The polarizer is a molecule that promotes sorting of PINs to the membrane section where it is most abundant (e.g. specific kinase that phosphorylates PIN). The regulator is a molecule that is activated by auxin and inhibits polarizer presence on the membranes (e.g. antagonizing phosphatase). Auxin application on a membranes section promotes regulator trafficking on that section, which in turn reduces the presence of the polarizer. Diffusion of the regulator over the surface of the cell results in concentrating the polarizer on the opposite cell side on which auxin was applied. Finally, the polarizer promotes PIN trafficking by tuning the auxin contribution parameter $IP_{mem}$ discussed in equation (14).

The model can be resumed by the following assumptions:

- Auxin active import across plasma membrane is sensed by the cell in order to determine the net auxin flow that promotes regulator binding to the membrane. Auxin import into the cell creates and auxin influx-efflux relation for each specific membrane section, which is sued to regulate the trafficking of the regulator species, and it is calculated as:

$$Grad_{mem} = \left( (I_{mem} - E_{mem}) + \sum_{i}^{m} \frac{(I_{mem_i} - E_{mem_i})}{distance(mem, mem_i)} \right) \frac{1000}{A_{cell}} \qquad (18)$$

(18) $Grad_{mem}$ is the auxin influx-efflux ratio specific to membrane section *mem*; $(I_{mem} - E_{mem})$ indicates the net amount of auxin crossing the membrane section *mem*; $distance(mem, mem_i)$ is the distance of membrane section $mem_i$ from our reference membrane section *mem*, calculated as the Euclidean distance between the two sections midpoints; $A_{cell}$ is the area of the cell. The metric is further normalized dividing by the cell area and amplified by a constant factor of 1000.

- Regulator and polarizer are expressed and degraded at a constant rate in the cytoplasm, while traffic to the membranes is promoted by auxin for both species, and modulated by the auxin gradient for the regulator:

$$\frac{dREG_{cell}}{dt} = b_{REG} - REG_{cell} \left( Kreg_{tr} + Kreg_{GradT} \frac{{Grad_{mem}}^4}{{Grad_{mem}}^4 + {Kreg_{GradK}}^4} \right) \frac{{IAA_{cell}}^2}{{IAA_{cell}}^2 + {Kreg_{IAA}}^2} - d_{REG}\, REG_{cell} \qquad (19)$$

$$\frac{dPOL_{cell}}{dt} = b_{POL} - POL_{cell}\, Kpol_{tr} \frac{{IAA_{cell}}^2}{{IAA_{cell}}^2 + {Kpol_{IAA}}^2} - d_{POL}\, POL_{cell} \qquad (20)$$

(19-20) $REG_{cell}$ and $POL_{cell}$ are the regulator and polarizer concentration inside the cell, respectively; $b_{REG}$ and $b_{POL}$ are the regulator and polarizer basal production rate, respectively; $Kreg_{tr}$ and $Kpol_{tr}$ are the regulator and polarizer trafficking rate, respectively; $Kreg_{GradT}$ and $Kreg_{GradK}$ are regulator maximum trafficking rate activation by auxin-gradient and half-max trafficking activation by auxin-gradient, respectively; $Grad_{mem}$ is the auxin-gradient on membrane section *mem; $IAA_{cell}$* is the amount of auxin inside the cell; $Kreg_{IAA}$ and $Kpol_{IAA}$ are regulator and polarizer half-max trafficking activation by auxin, respectively; $d_{REG}$ and $d_{POL}$ are the regulator and polarizer degradation rate, respectively.

- The presence of auxin inside the cell promotes trafficking of the two molecules to the membranes. The amount of molecules present on a specific membrane section *mem* is defined by the following equations:

$$\frac{dREG_{mem}}{dt} = REG_{cell} \left( Kreg_{tr} + Kreg_{GradT} \frac{{Grad_{mem}}^4}{{Grad_{mem}}^4 + {Kreg_{GradK}}^4} \right) \frac{L_{mem}}{\sum_i^m L_{mem_i}} \frac{{IAA_{cell}}^2}{{IAA_{cell}}^2 + {Kreg_{IAA}}^2} \qquad (21)$$

$$+D(REG_{mem}) - d_{REG}REG_{mem}$$

$$\frac{dPOL_{mem}}{dt} = \qquad (22)$$

$$POL_{cell}\,Kpol_{tr}\,\frac{L_{mem}}{\sum_i^m L_{mem}}\frac{IAA_{cell}^{\ 2}}{IAA_{cell}^{\ 2} + Kpol_{IAA}^{\ 2}} + D(POL_{mem}) + G(POL_{mem}) - d_{POL}POL_{mem}$$

(21-22) $REG_{mem}$ and $POL_{mem}$ the regulator and polarizer on membrane section *mem*, respectively; $Kreg_{tr}$ and $Kpol_{tr}$ are regulator and polarizer trafficking rates, respectively; $REG_{cell}$ and $POL_{cell}$ are cytoplasmic regulator and polarizer, respectively; $Kreg_{IAA}$ and $Kpol_{IAA}$ are regulator and polarizer half-max trafficking activation by auxin, respectively; $L_{mem}$ is the length of the membrane section *mem*; $IAA_{cell}$ is the amount of auxin inside the cell; $D(REG_{mem})$ and $D(POL_{mem})$ are the net amount of regulator and polarizer diffused to membrane section *mem*, respectively; $G(POL_{mem})$ is the net amount of polarizer displaced by the regulator on membrane section *mem*; $d_{REG}$ and $d_{POL}$ are regulator and polarizer decay rate, respectively; $Kreg_{GradT}$ and $Kreg_{GradK}$ are regulator maximum trafficking rate activation by auxin-gradient and half-max trafficking activation by auxin-gradient, respectively; $Grad_{mem}$ is the auxin-gradient on membrane section *mem*.

- Regulator and polarizer diffuses along the cell membrane according to the following equations:

$$D(REG_{mem}) = D_{REG}\sum_{i}^{mem\pm1}\left(REG_{mem_i} - REG_{mem}\right) \qquad (23)$$

$$D(POL_{mem}) = D_{POL}\sum_{i}^{mem\pm1}\left(POL_{mem_i} - POL_{mem}\right) \qquad (24)$$

(23-24) $D(REG_{mem})$ and $D(POL_{mem})$ are the regulator and polarizer diffused over the membrane, respectively; $D_{REG}$ and $D_{POL}$ are regulator and polarizer diffusion rate, respectively; $REG_{mem}$ and $POL_{mem}$ are the regulator and polarizer on membrane section *mem*, respectively; The polarizer is displaced by the presence of regulator molecules towards the zone where the concentration of the regular is the lowest. To achieve this, we apply a simple stochastic algorithm based on a previous model for designing of self-organizing cell polarity(Chau et al., 2012): for a given membrane section *mem*, a fixed batch amount of polarizer is reserved for possible displacement; then one of the two adjacent membrane segments is selected randomly; if the selected segment contains less amount of regulator than the current segment, the batch of polarizer is moved to that segment, otherwise, nothing is performed. Polarizer displacement can be resumed by the following formula:

$$G(POL_{mem}) = Kdisp_{POL}\ if(\ REG_{mem+i} > REG_{mem}\ then\ POL_{mem+i}\ else\ 0$$
$$i = random(-1, +1) \quad (25)$$

(25) $G(POL_{mem})$ is the amount of polarizer displaced by the regulator and $Kdisp_{POL}$ is the polarizer displacement rate.

- Given the calculated amount of polarizer on a given membrane, the auxin flow contribution parameter $IP_{mem}$ on PIN sensitivity described in equation (14):

$$IP_{mem} = \frac{{POL_{mem}}^4}{{POL_{mem}}^4 + {Kpol_{IP}}^4} \quad (26)$$

(26) $Kpol_{IP}$ is the half-max value of polarizer $POL_{mem}$ for auxin flow contribution on PIN sensitivity.

#### 3 Root Growth Model

The classical morphogen gradient model dictates that the cell fate is regulated by the positional information encoded in different morphogen levels at different positions across a static tissue(Wolpert, 1969). However, in a growing system, cells are displaced quickly along the tissue hence varying their exposure to morphogens concentration depending on their current distance for the morphogens source(s). Moreover, cell growth dilutes morphogens concentration and influences their effect on cell regulation. The current model tries to address these issues by monitoring the combined effect of cell growth and morphogens concentration on root development. The proposed model for root growth includes the following assumptions:

1. Cells expand according to the chemiosmotic theory of auxin transport(Rayle and Cleland, 1992). Cells are under constant turgor pressure which is resisted by the elastic effect of cell walls. In the current model osmotic pressure is realized by a specific position-based dynamics constraint described in the following section.
2. Increasing auxin concentration induces the relaxation of the cell walls allowing cell expansion under the effect of turgor pressure. On the contrary, high auxin levels disable cell walls relaxation(Fendrych et al., 2018). The relationship between cell walls stiffness and auxin is expressed by the following formula:

$$kE_{wall} = kE_{Max}\left(\frac{{K_{1\,IAA}}^4}{{IAA_{cell}}^4 + {K_{1IAA}}^4} + \frac{{IAA_{cell}}^4}{{IAA_{cell}}^4 + {K_{2IAA}}^4}\right) \quad (27)$$

(27) $kE_{wall}$ is the extensional stiffness of the cell wall; $kE_{Max}$ is the maximum stiffness a cell wall can achieve; $K_{1\,IAA}$ is the auxin-induced cell wall relaxation coefficient; $K_{2IAA}$ is the auxin-induced cell wall stiffening coefficient; $IAA_{cell}$ is the auxin concentration inside the cell.

3. Cell growth is directionally constrained by the action of cortical microtubules, which, depending on their orientation, make anisotropic growth possible(Baskin, 2005). During model simulation the action of cortical microtubules is made possible by the action of a specific strain-based dynamics constraint (see following section). For a given cell, the strain stiffness of a single wall segment depends on its alignment with cortical microtubules: the closest the alignment the harder the stiffness. This rule prevents the expansion of cell walls that are constrained by microfibrils.
4. Cell division occurs when a certain area threshold is reached. The cell division plane passes through the centroid of the cell polygon and it is orthogonal to the cortical microtubules. As already discussed before, microtubules orientation defines growth anisotropy, and therefore the division line is defined accordingly. Cells divide symmetrically into two cell of the same type of the mother. There are however exceptions to the previous rule, which are justified by experimental observation and tailored to certain conditions:
    a. Cortical/Endodermis Initials Daughters (CEID) cells always divide parallel to cell polarity, to generate one cell of the same type and one CEID cell(Miyashima and Nakajima, 2011; Mylona et al., 2002).
    b. Cortical/Endodermis Initials Daughters (CEID) divide asymmetrically to generate one endodermal cell and one cortical cell(Miyashima and Nakajima, 2011; Mylona et al., 2002).
    c. Epidermis/Lateral Root Cap initials alternatively divide either parallel and orthogonal to cell polarity in order to produce lateral root cap cells or epidermal cells, respectively(Kumpf and Nowack, 2015).
    d. Quiescent cells divide rarely(Della Rovere et al., 2016), and therefore for sake of simplicity we assume that they never grow or divide.
    e. Columella initials divide asymmetrically to generate one cell of the same type and one columella cell(Scheres et al., 2002).
    f. Vascular initials divide asymmetrically to generate one cell of the same type and one vascular cell(Baum et al., 2002).

#### 4 Physical-based Graphics Implementation

The typical approach to simulate dynamic growing systems in biology is based on force calculations(Nealen et al., 2006).Tissues are usually represented as triangulated meshes made of connected vertices and forces are accumulated on these vertices following specific biological criteria such as internal turgor pressure, anisotropic expansion or gravity. Vertex acceleration is later derived from these forces and vertex masses according to Newton’s second law. A time integration scheme is then used to first compute the velocities from the accelerations and then the final positions from the velocities. Classical integration methods are usually unstable or very computationally expensive, resulting in either unmanageable or extremely inefficient simulations. Therefore, instead of a forced-based system we decided to implement the mechanical growth of *Arabidopsis thaliana* root using Position-Based Dynamics (PBD)(Müller et al., 2007). PBD is a recent method used to simulate physical phenomena like cloth, deformation, fluids, fractures, rigidness and much more(Müller et al., 2007). PBD omits the velocity layer and immediately works on the positions following constraints that restrict the systems dynamics in order to obtain a desired effect. PBD main loop inside the model code can be summarized by the following diagram:

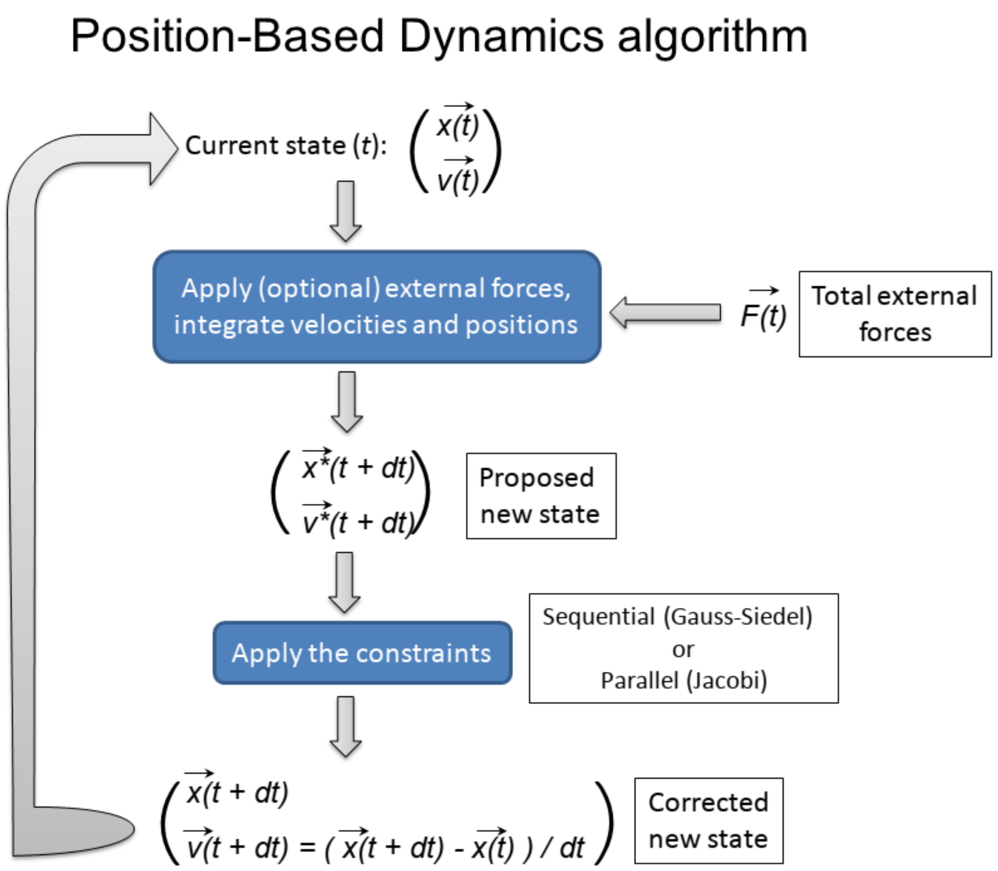

The generic algorithm for PBD is described as follow is pseudo-code:

```
// The algorithm assumes that each vertex in the model possesses the following
// attributes: velocity, position and mass.
// Vertices velocities are updated by applying the external forces (according to
// Newton's law) and it proposes new positions for each vertex.
for Vertex v in Vertices {
       v.previous_position = v.position;
       v.velocity += dt * (v.forces / v.mass);
       v.position += dt * v.velocity;
}

// The iterative solver applies the constraints to the proposed vertices
// positions, adjusting them such that they satisfy the constraints.
for i in iterations
       project_constraints(contraints, vertices);

// Finally, adjust vertices velocity based on the new adjusted positions
for Vertex v in Vertices
       v.velocity = (v.position - v.previous_position) / dt;
```

To provide a simple example of PBD constraint, consider the typical mass-spring system where two masses (usually represented by mesh vertices) are connected by an elastic spring. The elasticity of the spring applies a force on the masses which induces acceleration and velocity. In the PBD formulation, the same is achieved by the projecting the constraint $\boldsymbol{C}(\mathbf{p}_1, \mathbf{p}_2) = |\mathbf{p}_1 - \mathbf{p}_2| - \boldsymbol{d}$; $\mathbf{p}_1$ and $\mathbf{p}_2$ are the vertices positions and $\boldsymbol{d}$ is the spring resting

length. The resulting corrections $\Delta \mathbf{p}_i$ are subsequently weighted according to the inverse masses $\mathbf{w}_i = \mathbf{1/m}_i$. Finally, to account for the non-linear effect of the stiffness correction and to make it independent of the number of iterations, we multiply the stiffness correction by $k' = 1 - (1 - k)^{1/n}$, as described section 3.3 of the original paper(Müller et al., 2007).
PBD has been implemented in the current model by following the system described in the original paper(Müller et al., 2007).
The current model implements distance, shape, strain, bending and pressure constraints (Supplementary Fig. S2A-B), according to the following rules:

1. The distance constraint controls the distance between connected vertices(Müller et al., 2007). This constraint is useful to simulate elastic and plastic behavior. Cell edges can be either internal to the cell (representing the internal cytoskeleton and pectin matrix) or on the border (representing the cell walls). Edges have a different stiffness for extension and compression. Cells are generally regarded as incompressible objects so we assume maximum compression stiffness. Internal edges are meant to represent the cell pectin matrix and implemented as a viscoelastic material. Border edges represent the cell walls and are very stiff in order to prevent cell swelling from turgor pressure but can be relaxed through the action of auxin to allow cell growth.
2. The shape constraint(Müller et al., 2007) simulates the mechanical forces involved in the preservation of the cell structure. The shape constraint prevents cell deformation and collapse under external forces while allowing cell growth under internal pressure.
3. The strain constraint makes anisotropic growth possible(Müller et al., 2014). This constraint is applied on mesh triangles restricting deformation according to cortical microtubules orientation. Disordered microtubules organization results in isotropic growth, while ordered microtubules produce anisotropic growth.
4. The pressure constraint(Müller et al., 2007) mimics cell osmotic pressure. This constraint is always active and allows cell expansion, which can be opposed by cell wall stiffness and restricted in a certain direction by the strain constraint (according to cortical microtubules organization). This constraint is different from previously discussed constraints because it also implements a special version of Position-Based Dynamics, called XPBD(Macklin et al., 2016). Simple PBD does not consider time in its original implementation, which makes it inadequate for growing constraints like pressure, since it does not replicate a physical realistic behavior. The pressure constraint has therefore been implemented in XPBD, making it more time-compliant.
5. The bending constraint prevents cell walls angle to drift too far away from the resting condition(Müller et al., 2007), hence avoiding cell collapse, mimicking cytoskeleton robustness.

### 5 Parameters Sensitivity and Hypothesis Testing

The proposed model can be easily extended to address additional important biological questions and to perform specific hypothesis testing. To show the model's potential in tackling these matters we performed a simple parameters sensitivity test that also addresses core biological questions explained in the main text. The model was put to the test by varying specific core parameters, in this way we could both assess the robustness of the model to parameters change and show how to perform a hypothesis test using the current model.

In the results section we described how auxin transport direction emerges from the dynamic interaction between auxin influx and efflux carriers. According to our model, PINs trafficking to a specific membrane section of the cell is determined by the joint interaction between auxin flow and CMTs orientation. Put shortly, CMTs orientation regulates PINs polarity, while auxin flow discriminates between the two available poles. As described before in equation (14), auxin flow contribution to PINs trafficking is modulated by the variable $IP_{mem}$. This variable is calculated in two different ways depending on whether we are running the model with the auxin-flux method or the regulator-polarizer method, but the final contribution to membrane's PIN sensitivity is determined by its coefficient *kP.* Therefore, we simulated root growth by setting the coefficient *kP* (the default value in the wild-type simulation is 3) over a different range of values (Supplementary Fig. S10A). Despite the broad range of values tested, the model is able to reproduce realistically looking simulations, demonstrating the robustness of the model against this core parameter. However, at closer inspection it appears clear that when the contribution of auxin flow to PINs trafficking is completely ignored ($kP = 0$), auxin is notably reduced compared to the wild-type situation ($kP = 3$), and auxin barely reaches tissues far from the QC (after refluxing back from the tip) and internal tissues that are usually replenished through lateralization are almost deprived of auxin (cortex and endodermis) (Supplementary Fig. S10A). On the contrary, by setting higher values ($kP >= 5$) auxin flow is much stronger (Supplementary Fig. S10A). In this situation, the auxin-reflux loop induced by auxin lateralization from the epidermis into the cortex is massively increased, creating zones of auxin accumulation in the reflux zone, and also higher auxin concentrations in the pericycle (Supplementary Fig. S10A).

In the results section we further described how in the model cell elongation is driven by the action of auxin on cell walls. Auxin induces cell's walls relaxation according to equation (27). The relationship between auxin and cell stiffness is regulated by $K_{1auxin}$, the auxin-induced cell wall relaxation coefficient. This parameter regulates cell stiffness response on auxin concentration. Therefore, we simulated root growth by setting the coefficient $K_{1auxin}$ (the default value in the wild-type simulation is 0.05) over a different range of values (Supplementary Fig. S10B). It can be observed that the model is able to reproduce realistically looking simulations for a range of values close to the default wild-type setting, demonstrating the robustness of the model against this core parameter (Supplementary Fig. S10B). Low values of $K_{1auxin}$ do not seem to produce any visible variable for the default configuration, while higher values notably reduce root growth, to almost completely stopping it (Supplementary Fig. S10B). The latter results happen because even higher concentration of auxin is required to trigger cell's walls relaxation, and the auxin influx provided with the original settings is not enough to reach it. This uninvolved simulation shows how the model can be used to test auxin effect on cell growth. Future modifications of the current model could test for more elaborated situations; for example by connecting auxin concentration with the enzymatic processes involved in cell walls relaxation.

Taken together, these simulations demonstrate how the current model can be easily expanded to include additional elements from biological observations and how to perform hypothesis testing for different model alternatives.

**Supplementary Table S1.**

Model parameters description and value. Parameters that were obtained from previous works show a reference.

| Parameter | Description | Value | Unit |
|---|---|---|---|
| **Cell Polarization Model** | | | |
| $R_{MF}$ | CMTs reorientation rate | 0.02 | $h^{-1}$ |
| $d_{MF}$ | CMTs degradation rate | 0.01 | $h^{-1}$ |
| **Auxin transport Model** | | | |
| $b_{IAA}$ | basal auxin production rate | 0* | nM/h |
| $DI_{IAA}$ | auxin diffusion rate in the intercellular space | 1 | $\mu m^2/h$ |
| $P_{IAA}$ | membrane auxin permeability | 0.2(Grieneisen et al., 2007) | μm/h |
| $d_{IAAb}$ | basal auxin degradation rate | 0.0125(Perianez-Rodriguez et al., 2021) | nM/h |
| $d_{IAAMax}$ | maximum auxin degradation rate coefficient | 0.125 | $h^{-1}$ |
| $K_{IAAMax}$ | coefficient for half-max auxin degradation | 5 | nM |
| $K_{AUX1}$ | coefficient of auxin importing rate by AUX/LAX | 1 | μm/h |
| $K_{PIN}$ | coefficient of auxin export rate by PIN | 1.4(Mironova et al., 2010) | μm/h |
| $b_{AUX1}$ | AUX1/LAX basal expression | 1 | nM/h |
| $AUX1_{expr}$ | auxin-induced AUX1/LAX maximal expression | 30 | nM/h |
| $AUX1_{K}$ | auxin-induced AUX1/LAX half-max expression | 0.01 | nM |
| $AUX1_{tr}$ | AUX1/LAX trafficking rate | 1 | $h^{-1}$ |
| $AUX1_{Max}$ | maximum concentration of AUX1/LAX | 2 | nM |
| $AUX1_{MaxMem}$ | maximum concentration of AUX1/LAX on membrane sections | 15 | nM |
| $d_{AUX1}$ | AUX1/LAX degradation rate | 0.08 | $h^{-1}$ |
| $b_{PIN}$ | PIN basal expression | 0.2(Mironova et al., 2010) | nM/h |
| $PIN_{expr}$ | auxin-induced PIN maximal expression | 50 | nM/h |
| $PIN_{K}$ | auxin-induced PIN half-max expression | 0.05 | nM |
| $PIN_{tr}$ | PIN trafficking rate | 1 | $h^{-1}$ |
| $PIN_{Max}$ | maximum PIN concentration inside the cell | 2 | nM |
| $PIN_{MaxMem}$ | maximum concentration of PIN on membrane sections | 15 | nM |
| $d_{PIN}$ | PIN degradation rate | 0.08(Mironova et al., 2010) | $h^{-1}$ |
| $d_{PINmax}$ | maximum PIN degradation rate on membranes | 0.8 | $h^{-1}$ |
| $kMF$ | coefficient for CMTs orientation contribution to PIN sensitivity | 0** | - |
| $kP$ | coefficient for auxin flow contribution to PIN sensitivity | 3 | - |
| $kMFP$ | coefficient for interaction CMTs orientation + auxin flow contribution to PIN sensitivity | 3 | - |
| $kG$ | coefficient for cell geometry contribution to PIN sensitivity | 3 | - |
| $Kmf$ | half-max CMTs orientation contribution to PIN sensitivity | 0.5 | - |
| $Kgeom$ | half-max cell geometry contribution to PIN sensitivity | 0.5 | - |
| **Auxin-flux and regulator-polarizer methods** | | | |

| | | | |
|---|---|---|---|
| *Kflux* | auxin-flux half-max contribution on PIN sensitivity | 0.1 | nM µm |
| $b_{REG}$ | regulator basal expression | 10 | nM/h |
| $b_{POL}$ | polarizer basal expression | 10 | nM/h |
| $d_{POL}$ | regulator decay rate | 0.08 | $h^{-1}$ |
| $d_{REG}$ | polarizer decay rate | 0.08 | $h^{-1}$ |
| $Kreg_{tr}$ | regulator base trafficking rate | 1 | $h^{-1}$ |
| $Kpol_{tr}$ | polarizer base trafficking rate | 0.01 | $h^{-1}$ |
| $D_{reg}$ | regulator diffusion rate | 1 | $\mu m^2/h$ |
| $D_{pol}$ | polarizer diffusion rate | 0.1 | $\mu m^2/h$ |
| $Kdisp_{POL}$ | polarizer displacement rate | 10 | $h^{-1}$ |
| $Kreg_{IAA}$ | regulator auxin-induced half-max trafficking rate | 0.01 | nM |
| $Kpol_{IAA}$ | polarizer auxin-induced half-max trafficking rate | 0.01 | nM |
| $Kreg_{GradT}$ | regulator max trafficking rate activation by auxin gradient | 1 | nM/h |
| $Kreg_{GradK}$ | regulator auxin gradient-induced half-max trafficking rate | 1 | nM |
| $Kpol_{IP}$ | half-max value of polarizer contribution on PIN sensitivity | 0.1 | nM |

**Root Growth Model**

| | | | |
|---|---|---|---|
| $kE_{Max}$ | cell wall maximum stiffness | 1 | - |
| $K_{1auxin}$ | half-max cell wall relaxation coefficient by auxin | 0.05 | nM |
| $K_{2auxin}$ | half-max cell wall stiffening coefficient by auxin | 3 | nM |

* auxin basal expression is set to zero for the default wild type model. However, when local production in the QC is necessary, the value is set to 10.

** this parameter is set to 0 in the default model and included in the formulas only for completeness.

#### Supplementary Figures

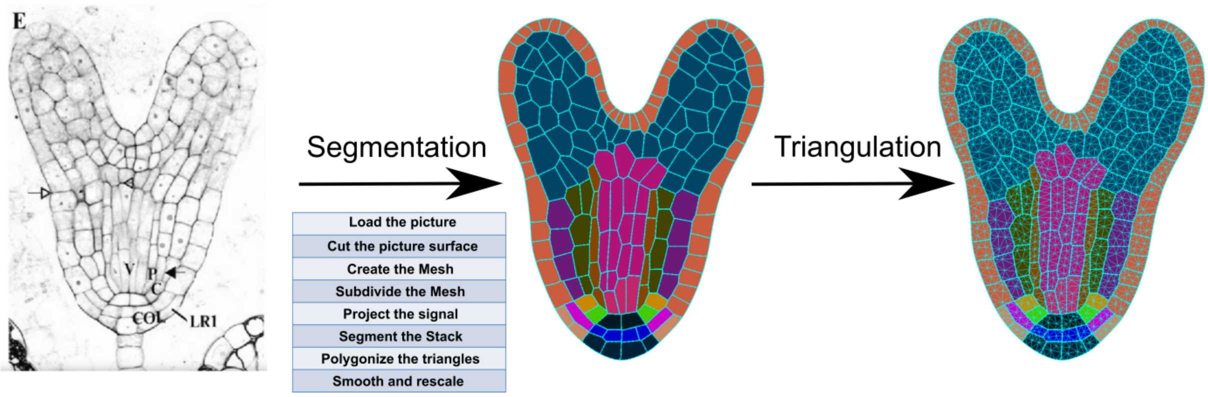

**Supplementary Fig. S1 Schematic procedure of the segmentation process that converts the microscopy picture of an A. thaliana embryo into a digitalized mesh.**

Microscopy pictures of *A. thaliana* embryo were initially loaded in the MorphoGraphX stack and the signal is projected against a digitalized mesh. Next, the image is segmented to recover the cell boundaries and the resulting mesh is triangulated to obtain an initial embryo template for the model.

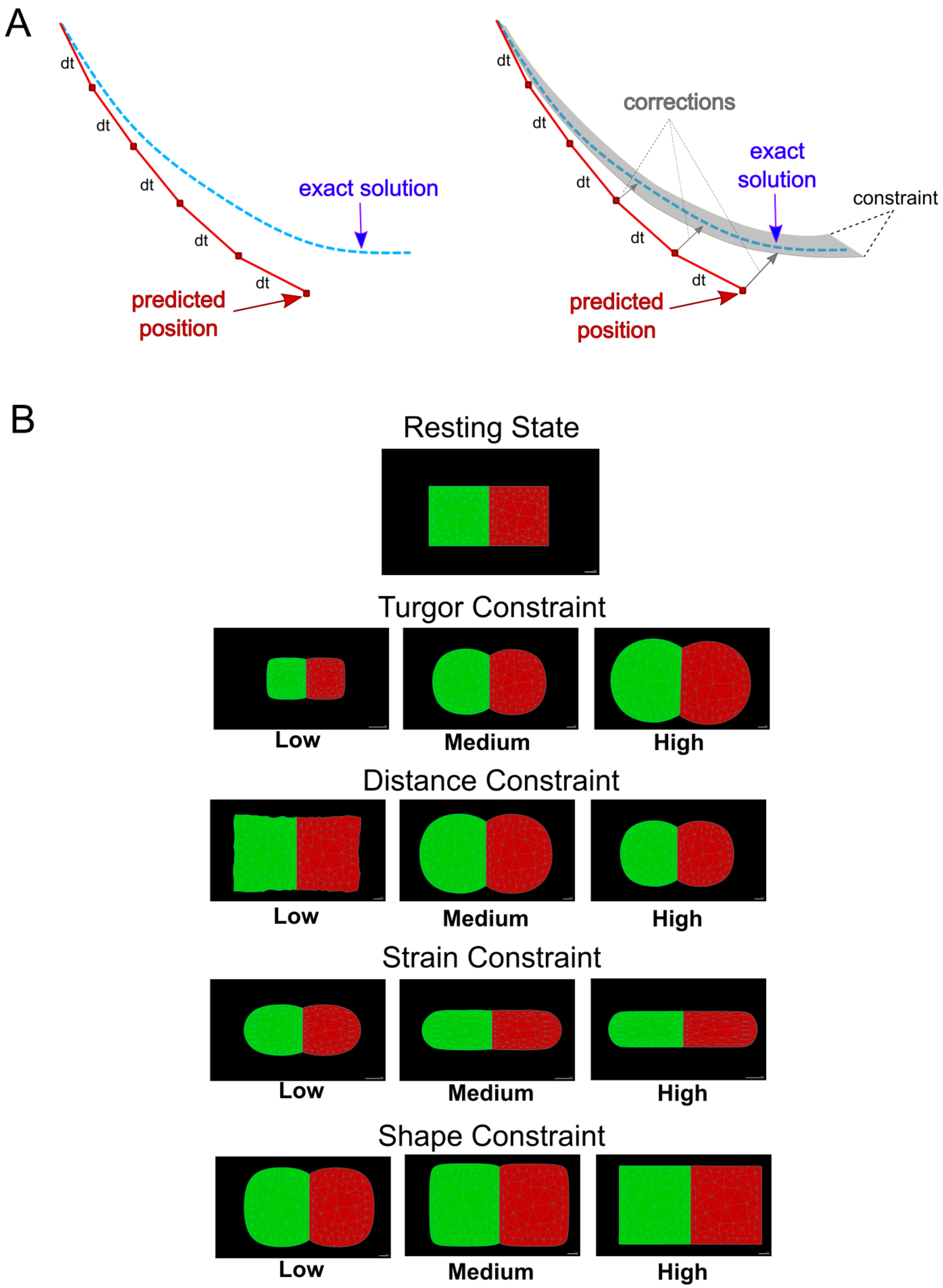

**Supplementary Fig. S2 Position-Based Dynamics implementation**

(**A**) Comparison between force-based methods (left panel) and position-based dynamics (right panel). In contrast to the typical force-based methods, position-based dynamics skips the velocity layer and directly works on objects positions in agreement with a determined set of constraints. (**B**) Schematic examples of PBD constraints. Starting from a simple template comprising two squared-shaped triangulated cells, simulations demonstrate the projected effect of each of the constraints implemented in the root model.

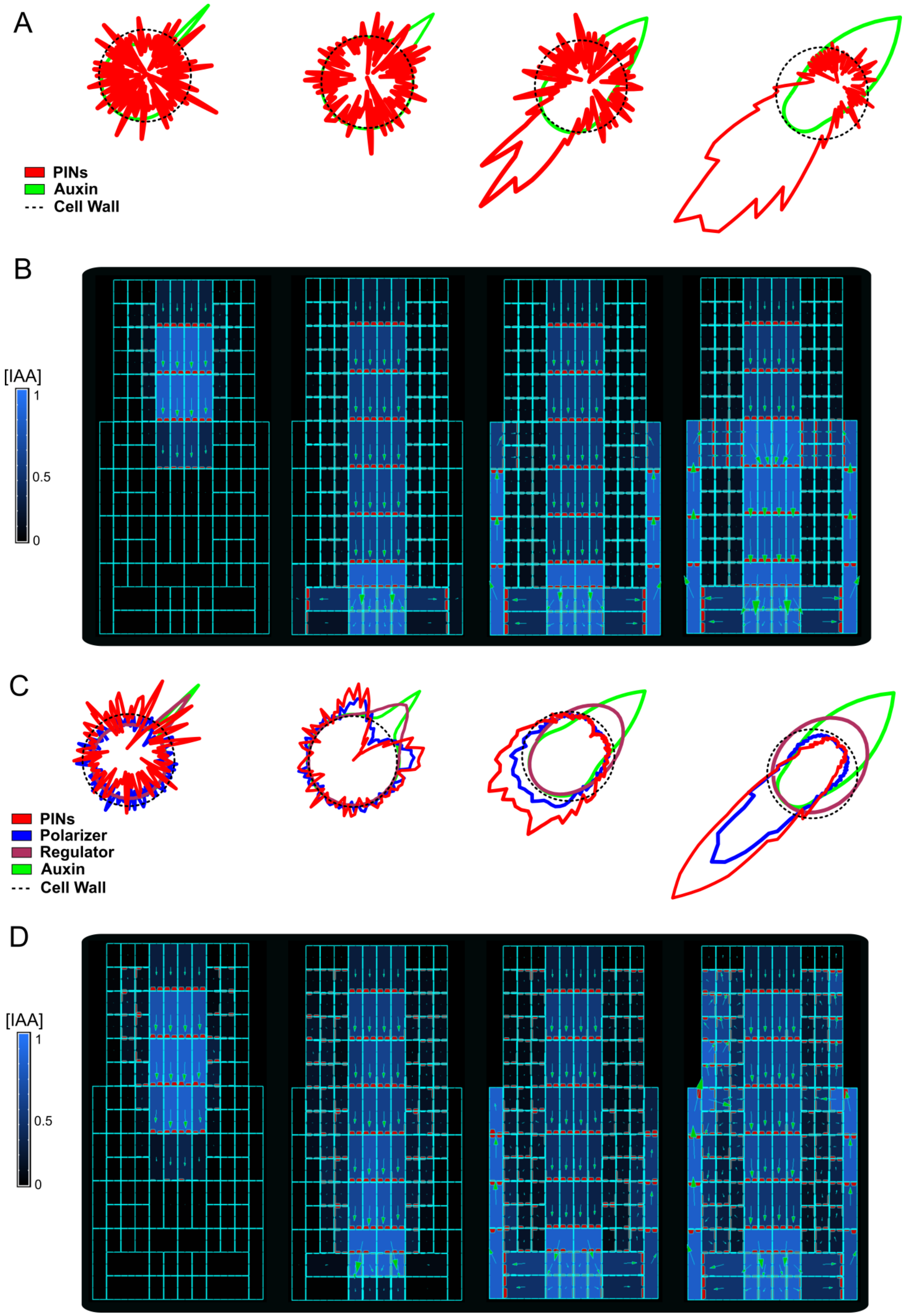
A
PINs
Auxin
Cell Wall
B
[IAA]
1
0.5
0
C
PINs
Polarizer
Regulator
Auxin
Cell Wall
D
[IAA]
1
0.5
0

935

**Supplementary Fig. S3 Comparison between auxin-flux and regulator-polarizer methods for PIN polarization.**

**(A)** Single-cell model simulation of the auxin-flux method. In this simple single cell model there are only two species of molecules: auxin (light green line) and PINs (red line). The auxin-flux method follows a very simple rule: auxin application on one side of the cell wall (dotted line) induces PINs polarization on the other side of the cell. **(B)** Model simulation using the auxin-flux method (A) on a simplified root-like grid structure. The auxin-flux method is capable of producing a realistic auxin flow inside the grid. Auxin was initially applied to the central cells of the top layer of the grid. (**C)** Single-cell model simulation of the regulator-polarizer method. This model represents an expansion of previous auxin-flux method (A). This model includes four molecular species: auxin (light green line), PINs (red line), a regulator (dark green line) and a PIN polarizer (blue line). Auxin application on one side of the cell wall (dotted line) induces recruitment of regulators proteins which inhibits the PIN polarizer pushing it towards the opposite side of the cell surface. The polarizer in turn fosters the recruitment of PINs molecules on that side of the cell. (**D)** Model simulation with the regulator-polarizer method (C) on a simplified root-like grid structure. The regulator-polarizer method is capable of producing a realistic and stochastic auxin flow inside the root. Auxin was initially applied to the central cells of the top layer of the grid.

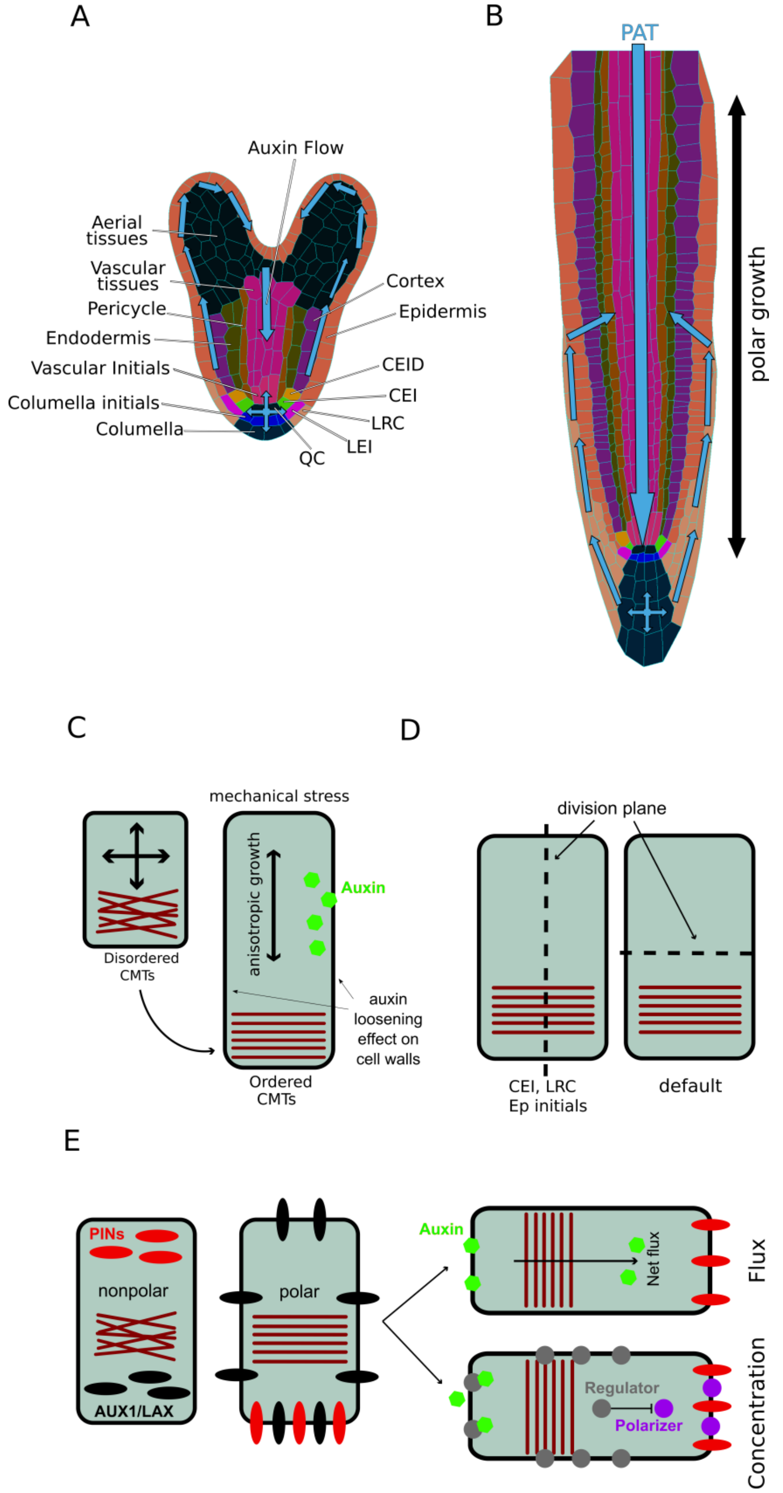

**Supplementary Fig. S4 Model purpose and design description.**

(**A**) Digitized microscopy image of an *A. thaliana* heart-stage embryo processed through MorphoGraphX(de Reuille et al., 2015). Cell types are shown with different colors, while estimated auxin flow is indicated with arrows. (**B**) Schematics of a mature *A. thaliana* root. The direction of auxin transport is visualized with blue arrows. Growth anisotropy is depicted with a bidirectional black arrow. (**C**) Cells are under constant internal turgor pressure. Auxin induces cell walls relaxation and allows cell expansion. External mechanical stress imposed on root cell walls forces the reorganization of the cortical microtubules (CMTs), which in turn generate the anisotropic growth. (**D**) Cell division occurs according to the cell polarity defined by CMTs orientation. Cell division plane is orthogonal to cell polarity; exceptions to this rule are cortex/endodermis and LRC/epidermis initials. (**E**) PINs localization is defined by CMTs orientation and auxin flux direction. Auxin feedback on PINs localization is determined by two equivalent mechanisms; the total auxin flux through plasma membrane (the auxin-flux method) or inhibition of PIN polarizer (e.g. kinase) through auxin-driven regulator (e.g. phosphatase) (the regulator-polarizer method).

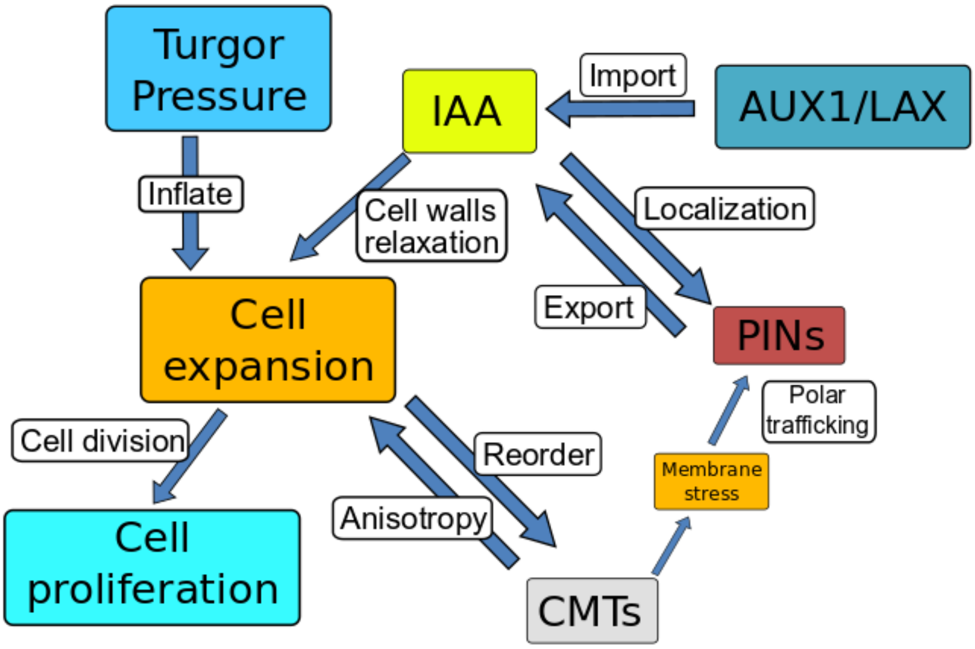

**Supplementary Fig. S5 Schematic diagram of the root model.**

The diagram shows the main components of the current model and their connections:

- Turgor pressure forces isotropic cell expansion driven by cell wall relaxation induced by auxin.
- Cell expansion causes mechanical stress on the cell walls that triggers cortical microtubules (CMTs) reorganization. Microtubules restrict cell expansion along the axis, causing anisotropic growth.
- Cells divide once the cell area to reach a specific threshold.
- CMTs alignment maintains cell polarity, reinforcing the auxin-dependent PINs polarization.
- Polar PINs efflux carriers and non-polar AUX1/LAX influx carriers mediate the auxin distribution along the root.

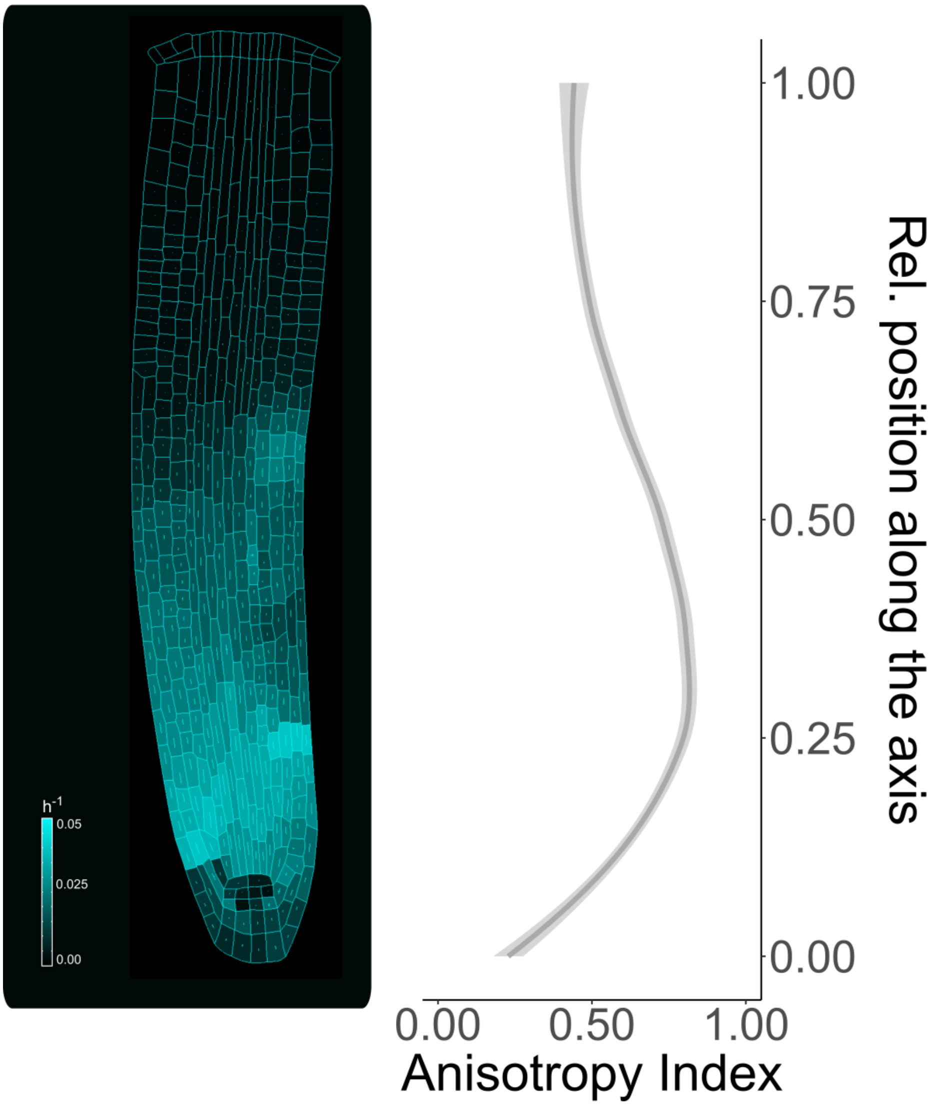

**Supplementary Fig. S6 Anisotropy index along the simulated root.**

Individual cells growth rate during model simulation (left figure) and anisotropy index along the root axis (right plot). The anisotropy index has been calculated as the difference between two orthogonal vectors representing the amount of cortical microtubules strength inside the cells. A high anisotropy index indicates a greater tendency to anisotropic growth, in contrast to a lower index which denotes a tendency towards the isotropic growth.

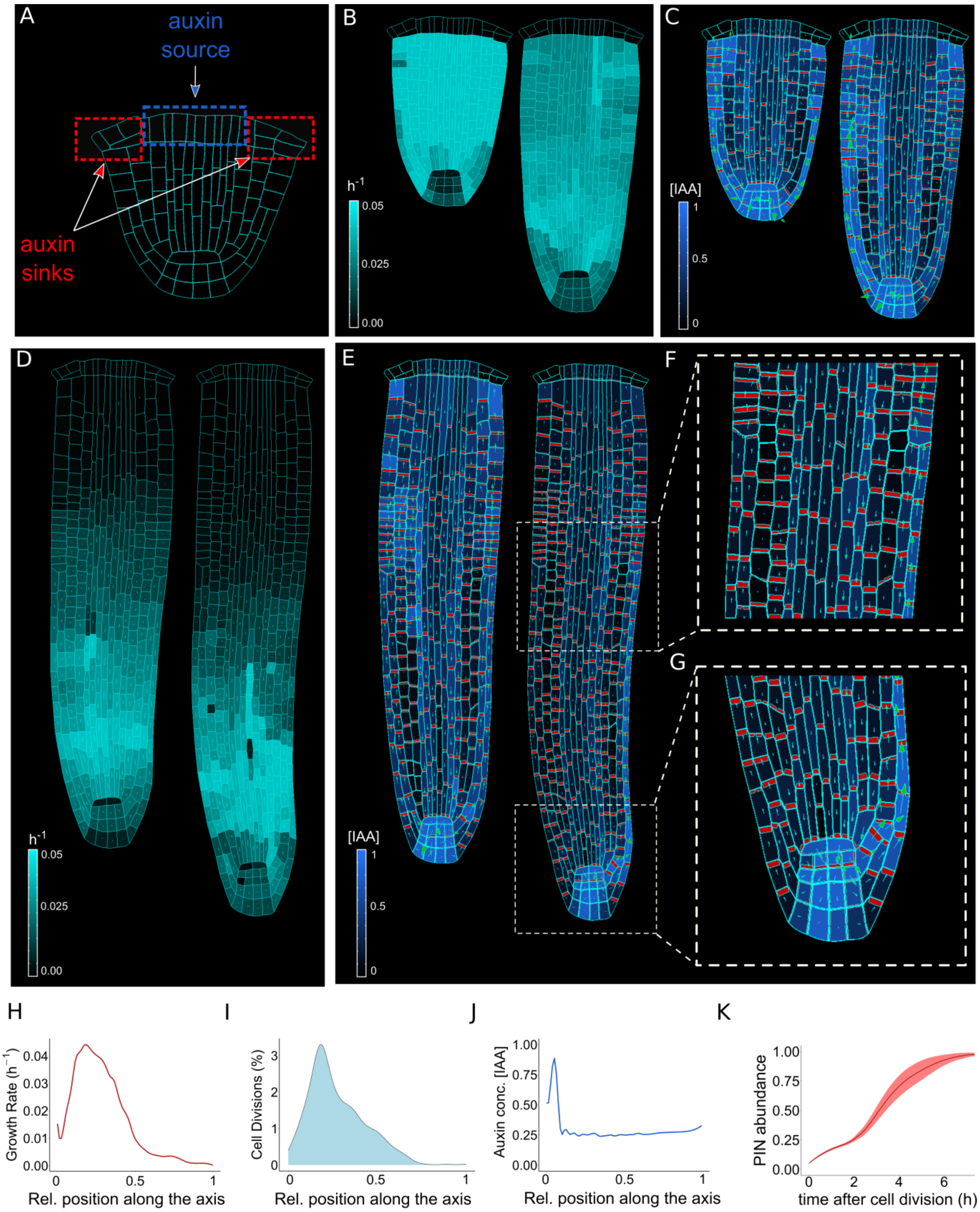

**Supplementary Fig. S7 The model is able to reproduce realistic root meristem geometry, auxin distribution and PINs localization. The simulations shown in this figure use the regulator-polarizer model.**

(**A**) Initial model template, indicating the location of auxin influx (auxin source) and evacuation (auxin sinks). (**B, D**) Time lapse of model simulation showing individual cells' growth rate (bright cyan color) and principal growth directions (white lines). (**C, E**) Time lapse of model simulation showing individual cells auxin concentration (blue color) and auxin flow (arrows).

(**F, G**) Model simulation close up on basal meristem (F) and root apical meristem (G). The model correctly reproduces experimentally observed PINs localization. Note the bipolar auxin flow in the cortical tissues (F). (**H**) Growth rate profile along the root axis. The fastest growing region is located in the apical meristem. (**I**) Cell division profile along the root axis. The majority of cell divisions occur in the apical meristem. (**J**) Auxin concentration profile along the root axis. Most of auxin is concentrated in the root tip. (**K**) Time lapse profile of PINs re-localization on the membranes after cell division. PINs re-localization is completed approximately 5 hours after cell division.

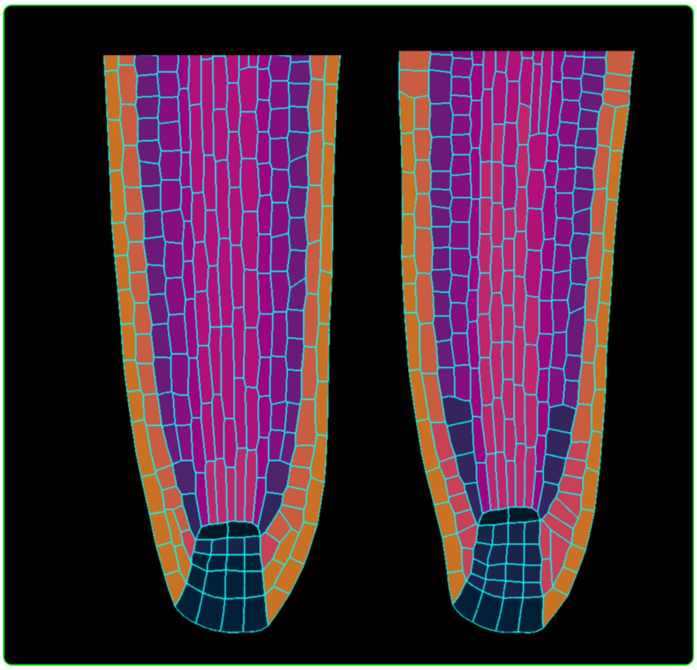

**Supplementary Fig. S8 Cell division rules testing**

Model testing of the cell division rule, comparing the wild-type simulation (left) with the simulation where specific rules for the stem cell niche are ignored (right).

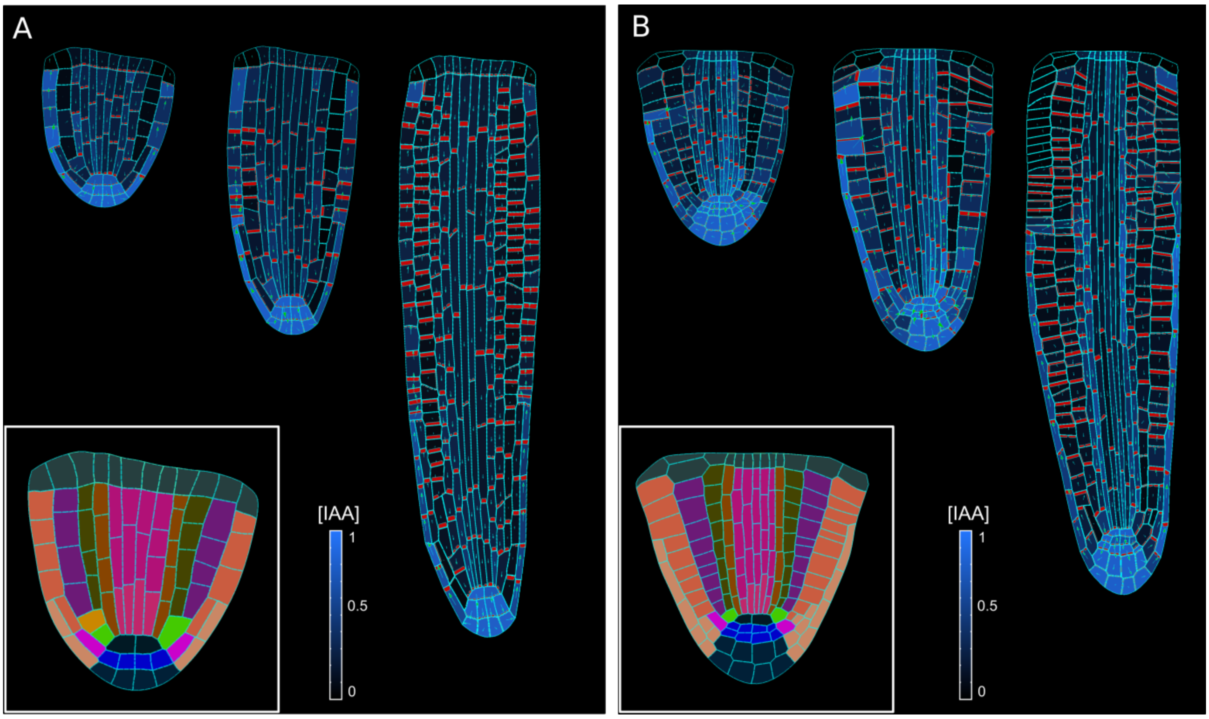

**Supplementary Fig. S9 Model simulations using alternative embryo templates.**

(**A,B**) Model simulations using alternative embryo templates(Nieuwland et al., 2016; Scheres et al., 1994). The particular choice of an initial embryo template has no impact on root self-organization of growth and auxin distribution.

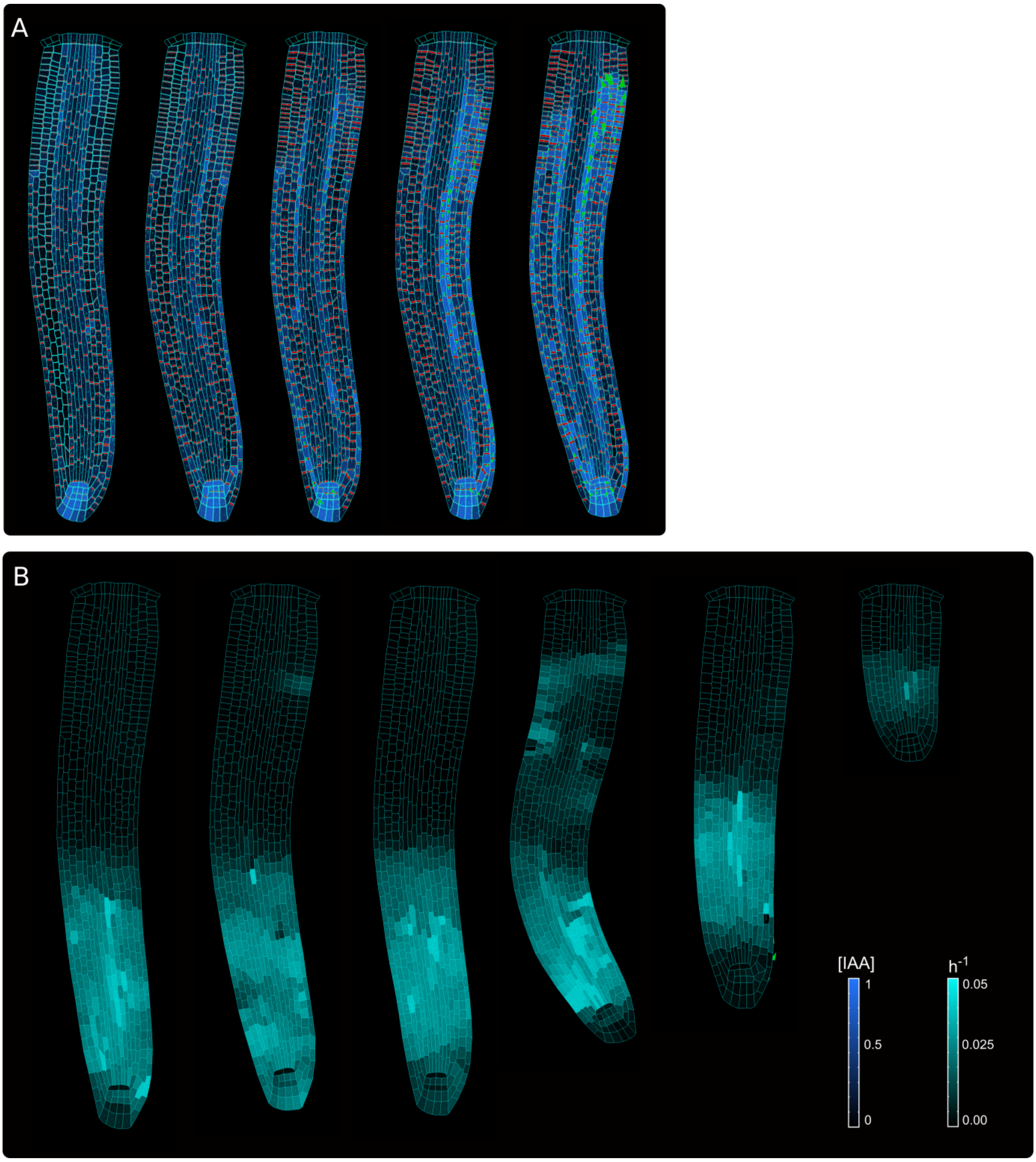

**Supplementary Fig. S10 Model parameters sensitivity**

**(A)** Model testing for parameter *kP,* the coefficient of contribution for auxin flow to PIN sensitivity for trafficking to the membranes (equation 14). The values tested are (for the left to right): 0, 1, 3 (default wild-type), 4 and 5. **(B)** Model testing for parameter $K_{1auxin}$, the auxin-induced cell wall relaxation coefficient (equation 27). The values tested are (for the left to right): 0.005, 0.01, 0.05 (default wild-type), 0.1 and 0.2.

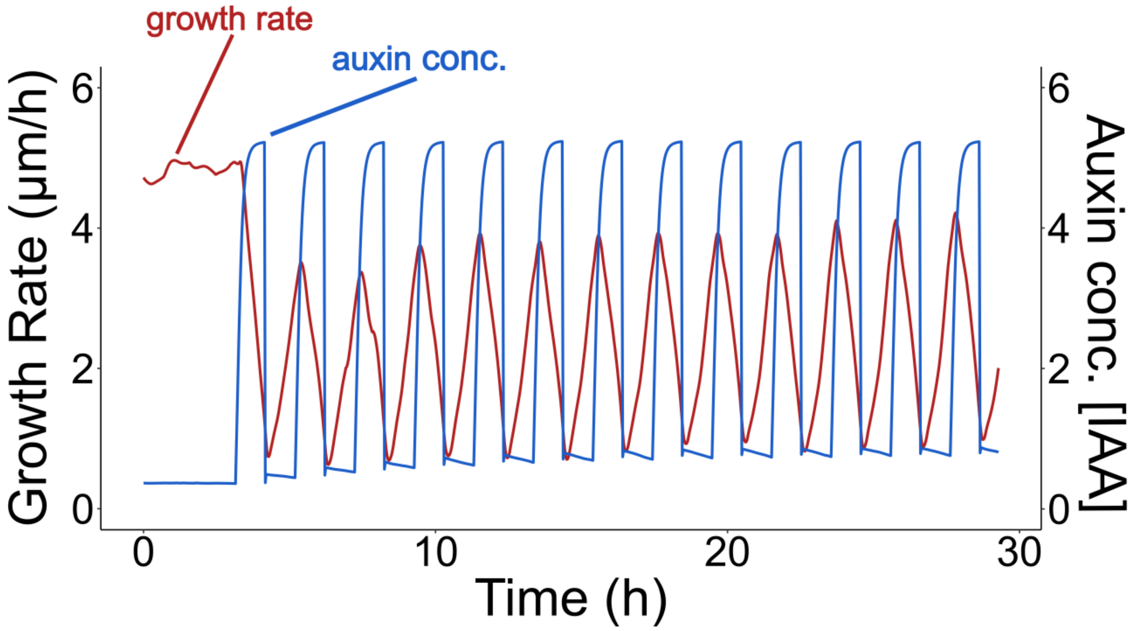

**Supplementary Fig. S11 Root growth reversible inhibition by external auxin application.**

Successive application of 10 nM of external auxin in model simulations according to a predefined cycle of 1 hour. Root growth is inhibited by the introduction of high amounts of auxin, and subsequently restored after the excess of auxin is removed.

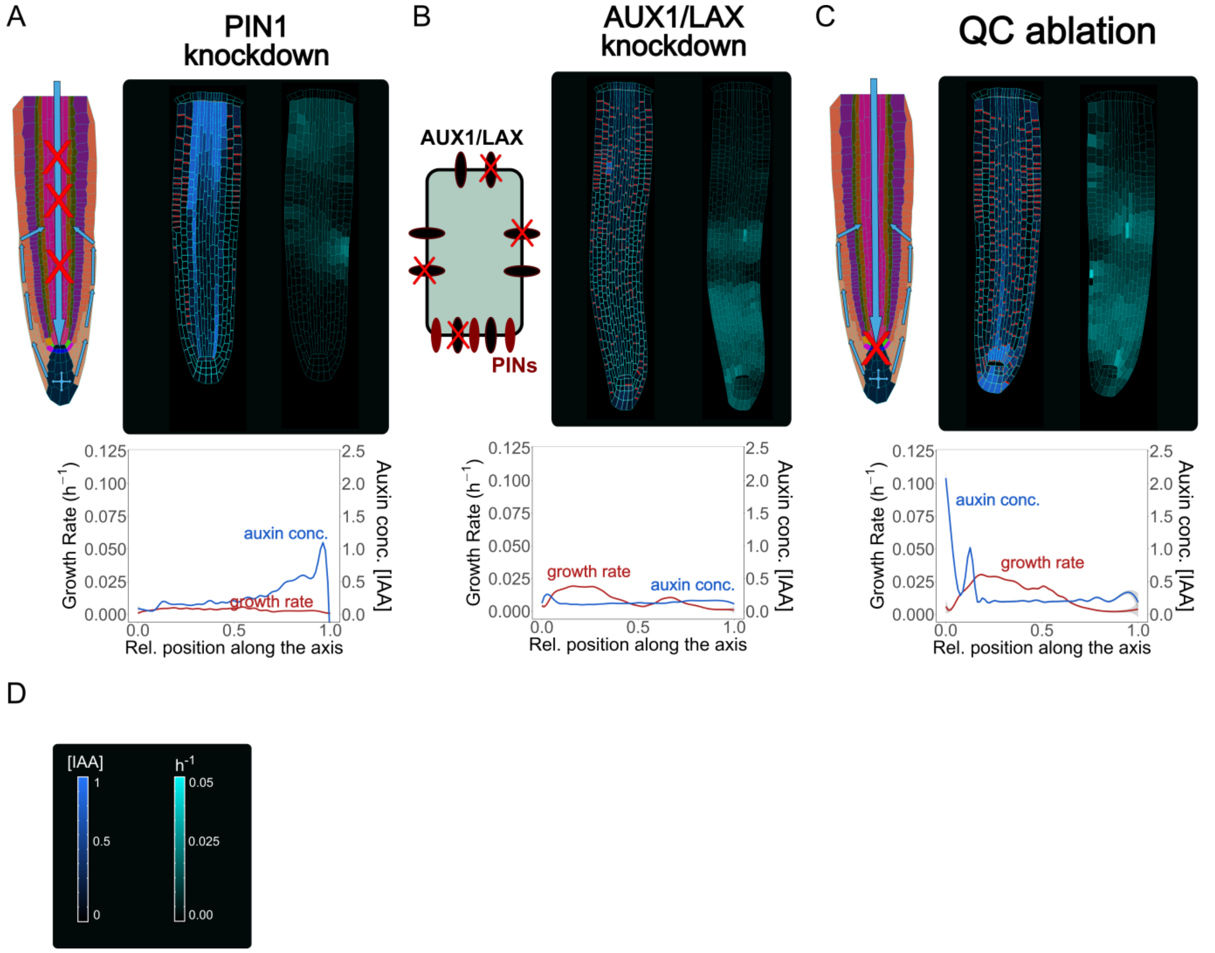

**Supplementary Fig. S12 Model simulations recapitulate experimentally observed phenotypes.**

(**A**) Model simulation of the *pin1* knockdown mutant. To replicate this type of mutation, PINs expression was reduced by 90% in the vascular tissues, pericycle and endodermis. As it can be observed, auxin levels in the root are strongly depleted as the hormone flow is severely affected by PIN1 down-regulation. (**B**) Simulated *aux1* knockdown mutant. AUX1/LAX expression was reduced by 90% in every cell. This mutant present a striking phenotype as auxin level and root growth are both significantly decreased. (**C**) Model simulation of QC ablation. Removing the QC cells results in an increase of auxin level in the vascular initial cells just above the ablated QC. The model also experiences the disappearance of acropetal auxin flow, as less auxin is able to reach and accumulate in the root tip. (**D**) Scale bars of auxin concentration and cell growth rate.

### Supplementary Video Legends

**Supplementary Video S1.**

Differential cell growth at the embryo RSI produces anisotropic expansion of the root, related to Fig. 1. The left section simulates root growth with uniform growth at RSI. The root is unable to achieve independent anisotropic growth since the whole embryo expands isotropically. The right section simulates root outgrowth by assuming differential growth rate at the RSI. Under these conditions the anisotropic growth of the root is self-organizing, later setting the polarity axis.

**Supplementary Video S2.**

Model simulations obtained with the auxin-flux method, related to Fig. 2. The upper simulation displays auxin distribution (blue color), auxin flow direction (arrows) and PIN localizations (red). The lower simulation displays cell growth rate (bright cyan color) and principal growth directions (white lines).

**Supplementary Video S3.**

Model simulations obtained with the regulator-polarizer method, related to Supplementary Fig. S7. The upper simulation displays auxin distribution (blue color), auxin flow direction (arrows) and PIN localizations (red). The lower simulation displays cell growth rate (bright cyan color) and principal growth directions (white lines).

**Supplementary Video S4.**

Model simulations with the enabled/disabled auxin reflux-loop and with/without local auxin production in the QC, related to Fig. 3. Only in reflux scenario auxin is permitted to lateralize from the epidermis back into the vascular tissues, which, together with a secondary auxin source in the QC is able to sustain the long-term root growth even after the removal of the main auxin source from the shoot.

**Supplementary Video S5.**

Successive application of external auxin in model simulations according to a predefined cycle, related to Fig. 4. Root growth is inhibited by the introduction of high amounts of auxin, and subsequently restored after the external application is stopped. The simulation displays auxin distribution (blue color), auxin flow direction (arrows) and PIN localizations (red).

**Supplementary Video S6.**

Model simulation of the *pin2* knockdown mutant, related to Fig. 5B. In-silico *pin2* mutant shows strongly reduced PINs expression in the lateral root cap, epidermis and cortex. Note that acropetal auxin flow is severely affected and auxin tends to accumulate in the lateral tissues.

**Supplementary Video S7.**

Model simulation of the *pin1* knockdown mutant, related to Supplementary Fig. S12A. To replicate this type of mutation, PINs expression was reduced by 90% in the vascular tissues, pericycle and endodermis. As it can be observed, auxin levels in the root are strongly depleted as the hormone flow is severely affected by PIN1 down-regulation.

**Supplementary Video S8.**

Model simulation of the *aux1* knockdown mutant, related to Supplementary Fig. S12B. AUX1/LAX expression was reduced by 90% in every cell. This mutant present a striking phenotype as auxin level and root growth are both significantly decreased.

**Supplementary Video S9.**

Model simulation of QC ablation, related to Supplementary Fig. S12C. Removing the QC cells results in an increase of auxin level in the vascular initial cells just above the ablated QC. The model also experiences the disappearance of acropetal auxin flow, as less auxin is able to reach and accumulate in the root tip.

**Supplementary Video S10.**

Model simulation of lateral root cap ablation, related to Fig. 5C. Mechanical removal of LRC resulted in the strong accumulation of auxin inside the root tip, largely because auxin cannot flow anymore shootward through outermost tissues whereas growth rate was not significantly affected.

**Supplementary Video S11.**

Model simulation of root tip cutting, related to Fig. 5D. Removing the root tip results in a general increase of auxin level in the central vascular tissues, as a result of the disappearance of acropetal auxin flow.

**Supplementary Video S12.**

Model simulation of CMTs disruption on root growth and polarity, related to Fig. 5E. CMTs disruption was simulated by inducing a fast degradation of cortical microtubules. Cells lose polarity and growth anisotropy, causing the root to expand and bulge radially.
